# Growth hormone remodels the 3D-structure of the mitochondria of inflammatory macrophages and promotes metabolic reprogramming

**DOI:** 10.1101/2022.08.22.504729

**Authors:** Blanca Soler Palacios, Ricardo Villares, Pilar Lucas, José Miguel-Rodríguez-Frade, Ana Cayuela, Jonathan G Piccirillo, Manuel Lombardía, David Delgado Gestoso, Cristina Risco, Fernando Corrales, Carlos Oscar S. Sorzano, Nuria Martínez, José Javier Conesa, Francisco J. Iborra, Mario Mellado

**Affiliations:** Department of Immunology and Oncology, National Center for Biotechnology/CSIC, Cantoblanco Campus, 28049-Madrid, Spain; Biocomputing Unit, National Center for Biotechnology/CSIC, Cantoblanco Campus, 28049-Madrid, Spain; Department of Macromolecular Structures, National Center for Biotechnology/CSIC), Cantoblanco Campus, 28049-Madrid, Spain; Functional Proteomics Laboratory, National Center for Biotechnology/CSIC, Cantoblanco Campus, 28049-Madrid, Spain; Cell Architecture & Organogenesis Program, Centro de Biologia Molecular Severo Ochoa, CSIC-UAM, Cantoblanco Campus, 28049 Madrid, Spain; Program for Systems Biology of Molecular Interactions and Regulation, Institute for Integrative Systems Biology (I2SysBio), Campus Burjassot/Paterna Parc Cientific, 46980, Valencia, Spain

## Abstract

Macrophages are a heterogeneous population of innate immune cells that support tissue homeostasis through their involvement in tissue development and repair, and pathogen defense. Emerging data reveal that metabolism may control macrophage polarization and function and, conversely, phenotypic polarization may drive metabolic reprogramming. Here, using biochemical analysis, correlative cryogenic fluorescence microscopy and cryo-focused ion-beam scanning electron microscopy, we demonstrate that growth hormone (GH) functions as a metabolic modulator to reprogram inflammatory GM-CSF-primed monocyte-derived macrophages (GM-MØ). We found that exogenous treatment of GM-MØ with recombinant human GH suppressed glycolysis, lactate production and non-mitochondrial respiration, and enhanced mitochondrial oxidative phosphorylation. Likewise, GH treatment augmented mitochondrial volume and altered mitochondrial dynamics, including the remodeling of the inner membrane to increase the density of cristae. Our data demonstrate that GH likely serves a modulatory role in the metabolism of inflammatory macrophages and suggest that metabolic reprogramming of macrophages should be considered a new target to intervene in multiple inflammatory diseases.

## Introduction

Macrophages are cells of innate immunity with a high capacity for clearing pathogens and cell debris through phagocytosis, and they also play increasingly defined roles in orchestrating tissue repair. To achieve all this, macrophages display high functional heterogeneity and plasticity. Indeed, depending on their tissue environment and on the activation of specific signaling pathways, they can exhibit pro- or anti-inflammatory functions and phenotypes (Gordon & Taylor, 2005).

Macrophages also display distinct energy metabolic profiles that are linked to their inflammatory status. For example, lipopolysaccharide/interferon (LPS/IFN)-activated M1 macrophages are characterized by enhanced glycolysis (Freemerman et al., 2014) and pentose phosphate pathway activity (Jha et al., 2015) and by suppressed oxidative phosphorylation (OxPhos) efficiency (Feingold et al., 2012), which is suited to the production of pro-inflammatory cytokines, reactive oxygen species (ROS), and nitric oxide (NO). By contrast IL-4-induced M2 macrophages have an efficient OxPhos system (Vats et al., 2006) and poor pentose phosphate pathway activity (Haschemi et al., 2012), together with enhanced arginase-1 expression and suppressed production of NO and ROS, facilitating the release of anti-inflammatory cytokines, growth factors, and polyamines (De Santa et al., 2019). Furthermore, alveolar macrophages, the most abundant immune cells in the lung in homeostasis, use glutamine as a principal substrate to fuel mitochondrial metabolism (Wang et al., 2020), and tumor-associated macrophages shift toward OxPhos and fatty acid oxidation and exhibit functions largely similar to M2 macrophages in a poor glucose environment to preserve their immunosuppressive roles (Puthenveetil & Dubey, 2020). Indeed, in the hypoxic microenvironment of solid tumors, tumor-associated macrophages enhance glycolysis via mTOR activation, reducing endothelial glucose availability (Wenes et al., 2016).

Growth hormone (GH) is produced and secreted by somatotrophic cells. Although originally implicated in somatic growth control, GH displays numerous functions (Ogle et al., 1992; Veldhuis et al., 2000; Velloso, 2008) including regulation of the immune system (Kimata & Fujimoto, 1994; Lu et al., 2010; W. J. Murphy et al., 1992; Welniak et al., 2002). Its receptor is expressed by many leukocyte subsets, and GH binding influences the function of B- and T-cells, natural killer cells and macrophages (Lu et al., 2010). For instance, recombinant human GH (rhGH) administration alters tolerization mechanisms in mice through activation of regulatory T-cells and modulation of Th17 cell plasticity. It also curtails the development of type I diabetes (Villares et al., 2013), and contributes to ameliorate collagen-induced arthritis symptoms (Villares et al., 2018). In human inflammatory diseases, rhGH administration limits mucosal inflammation in experimental colitis (Han et al., 2007), and has protective effects in patients with active Crohn’s disease (Slonim AE et al., 2000). In myeloid cells, GH functions as a macrophage-activating factor (Warwick-Davies et al., 1995), and stimulates the proliferation of RAW 264.7 macrophages (Smith et al., 2000). It also plays an important role in granulocyte-macrophage colony-stimulating factor (GM-CSF)-derived macrophage (GM-MØ) reprograming towards an anti-inflammatory and reparative profile both *in vitro* and *in vivo* (Soler Palacios et al., 2020). Finally, by downregulating the NLRP3 inflammasome in macrophages, GH has been linked to pro-longevity effects that maintain immune system homeostasis during aging (Spadaro et al., 2016).

We report here that rhGH treatment reduces glycolysis, lactate production, non-mitochondrial respiration and ATP generation in inflammatory macrophages *in vitro*, and modifies their bioenergetic profile towards OxPhos. These findings were confirmed by examining the effect of rhGH on several important metabolic enzymes, including pyruvate kinase M2 (PKM2), ATP-citrate synthase (ACLY) and lactate dehydrogenase A (LDHA), which were all downregulated in rhGH-treated cells when compared with untreated cells. We also found that rhGH treatment of GM-MØ promoted a reduction in the levels of itaconate, an essential metabolite involved in immunomodulation of inflammatory macrophages (Hooftman et al., 2020). A more detailed analysis demonstrated that rhGH treatment of GM-MØ additionally affected the total mitochondria mass by promoting fusion and reducing mitophagy. These results were validated using confocal live-cell imaging combined with correlative cryogenic fluorescence microscopy and cryo-focused ion-beam scanning electron microscopy (cryo-FIB-SEM). Finally, our results indicated a rhGH-mediated increase of GM-MØ mitochondrial volume and remodeling of the inner mitochondria membranes to increase the density of cristae.

Altogether, these data indicate that GH acts as a metabolic modulator of GM-MØ, and modifies the mass, dynamics and internal structure of their mitochondria. Our findings also underscore the importance of cellular metabolism in the coordination of the immune response to environmental conditions and suggest new targets to treat diseases involving macrophages.

## Results

### Growth hormone reprograms human GM-MØ metabolism by suppressing glycolysis

It was previously demonstrated that the oxygen consumption rate (OCR) and aerobic glycolysis (measured as the extracellular acidification rate [ECAR]) are higher in GM-MØ than in M-CSF-derived macrophages (M-MØ), concomitant with elevated expression of genes encoding glycolytic enzymes (Izquierdo et al., 2015). However, M-MØ exhibited a significantly higher OCR/ECAR ratio (Izquierdo et al., 2015) and showed differences in iron and folate metabolism (Sierra-Filardi et al., 2014). These findings point to a potentially important role for metabolic reprogramming in macrophage polarization and function.

We previously reported that rhGH-stimulated GM-MØ have an anti-inflammatory and reparative profile (Soler Palacios et al., 2020), and we hypothesized that, under such experimental conditions, GH might influence inflammatory macrophage metabolism and reprogramming. To test this, we first performed gene-set enrichment analysis (GSEA) of RNA-sequencing data from rhGH-treated GM-MØ (Soler Palacios et al., 2020), observing a significant downregulation of the gene set involved in glycolysis compared with the untreated cells (Fig. 1A). We next surveyed the bioenergetic profile of GM-MØ and M-MØ, untreated or treated with rhGH, on the Seahorse XF-analyzer platform. Results showed that the OCR and ECAR metrics were higher in GM-MØ than in M-MØ, and both metrics were reduced in rhGH-treated GM-MØ (Fig. 1B–E). Additionally, rhGH treatment of GM-MØ changed their mitochondrial bioenergetics profile, as evidenced by a significant increase in the oxidative/glycolytic parameter ratio (Fig. 1F). By contrast, rhGH treatment failed to significantly affect M-MØ metabolism (Fig. 1). Overall, these data indicate that rhGH changes the balance between the glycolytic and respiratory activities of inflammatory macrophages and suggest that rhGH triggers metabolic reprogramming towards an anti-inflammatory phenotype.

**Figure 1.**
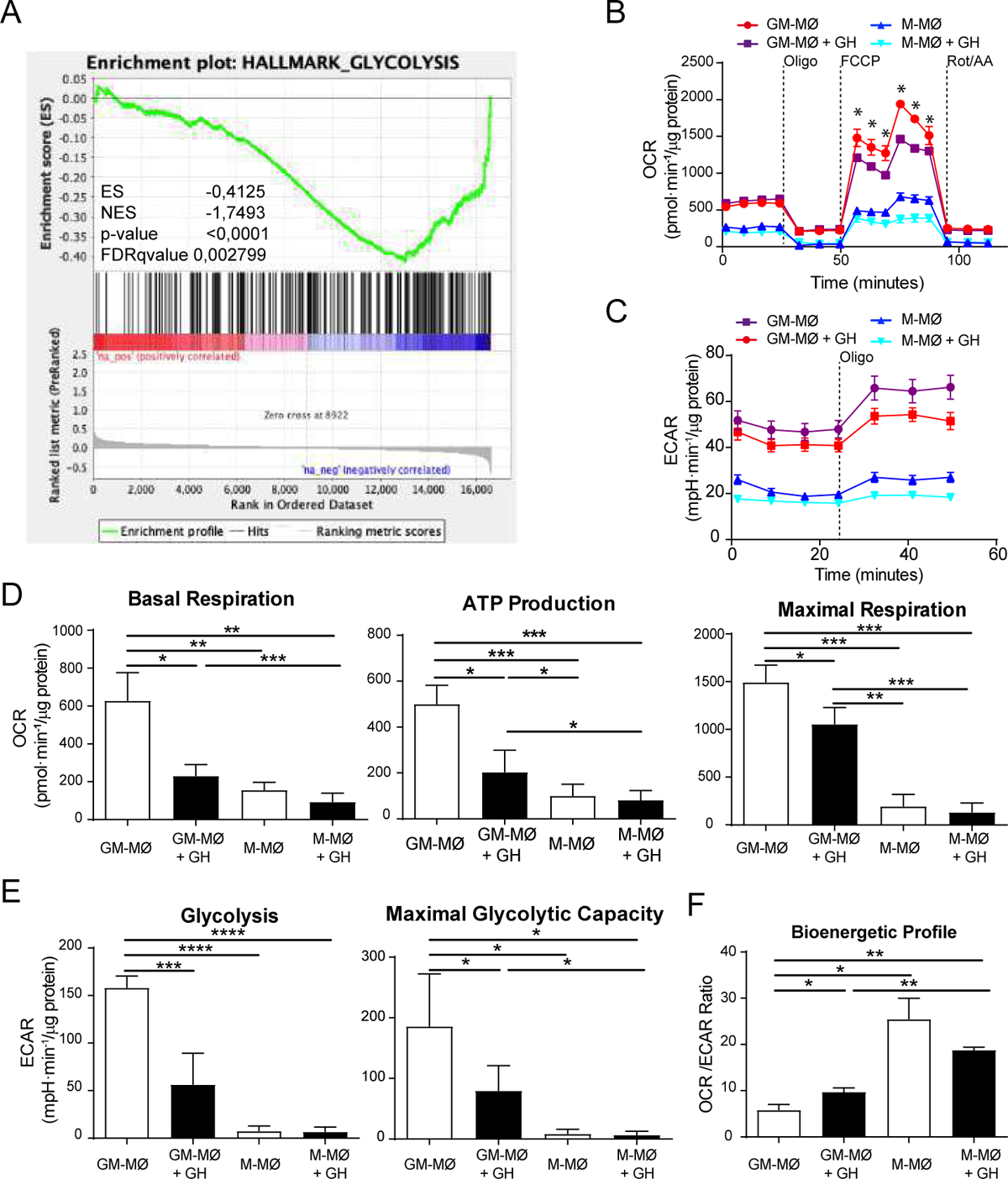
Treatment with rhGH suppresses glycolysis in inflammatory macrophages. **(A)** GSEA on the ‘log fold change-ranked’ list of genes obtained from untreated *versus* rhGH-treated GM-MØ according to limma analysis, using the previously defined glycolysis genes set (Glycolysis Hallmark). Vertical black lines indicate the position of each of the genes comprising the glycolysis gene-set. **(B)** OCR and **(C)** ECAR profiles of GM-MØ and M-MØ treated or not with rhGH (1 µg/ml, 24 h. 37°C) monitored using the Seahorse Bioscience extracellular flux analyzer. For the OCR profile, cells were sequentially treated, as indicated, with 1 µM oligomycin (Oligo), 0.6 and 0.4 μM FCCP and 1 μM rotenone plus 1 μM antimycin (Rot/AA). For the ECAR profile, measurements were performed in steady-state and after treatment with 1 μM oligomycin (Oligo). **(D)** Metabolic parameters obtained from the OCR profile after subtraction of the rotenone/antimycin-insensitive respiration. Results were normalized according the protein concentration and showed as mean ± SD (n=8). Paired *t*-test *p<0.05, **p<0.01, ***p<0.001. **(E)** Metabolic parameters obtained from ECAR profiling. Basal refers to the glycolysis rates in the absence of effectors, and maximal glycolytic capacity corresponds to ECAR value after inhibition of mitochondrial ATP synthesis with oligomycin. Results were normalized and shown as in (D). Results represent mean ± SD (n=8) Paired *t*-test *p<0.05, ****p<0.0001. **(F)** Bioenergetic profile (OCR/ECAR ratio) of GM-MØ and M-MØ under basal conditions and after stimulation with rhGH (1 μg/ml, 24 h, 37°C). Results are presented as mean ± SD (n=8). Results represent mean ± SD (n=8) Paired *t*-test *p<0.05, **p<0.01.

### Growth hormone downregulates the expression of key enzymes involved in glucose metabolism in GM-MØ

We next questioned whether the evident metabolic reprogramming of GM-MØ by GH correlated with differences in the expression of several key enzymes involved in aerobic glycolysis. We thus examined the expression of PKM2 (pyruvate kinase M2), and LDHA (lactate dehydrogenase A) in untreated or rhGH-treated GM-MØ fixed and immunostained with specific antibodies (Supplemental Fig. 1A). Untreated M-MØ were also included as a control. Quantification of the mean fluorescence levels of the images revealed that the expression of both proteins was significantly lower in rhGH-treated GM-MØ than in equivalent untreated cells (Fig 2A, B). In addition, qRT-PCR analysis of the same cells demonstrated that the gene expression levels of the glucose transporter *GLUT1* and *LDHA* were significantly lower after rhGH treatment (Fig. 2C). These findings suggest that GH promotes an increase in the amount of pyruvate in the mitochondria, which would increase the flux of acetyl-CoA through the tricarboxylic acid (TCA) cycle, and increase the production of NADH and subsequent O_2_ consumption as the final electron acceptor in the electron transport chain. Supporting this, we detected a significant decrease in lactate accumulation (Fig. 2D) and ROS production (Fig. 2E, F; Supplemental Fig. 1A, B) in GM-MØ treated with rhGH. Altogether, these data indicate that rhGH tempers aerobic glycolysis in GM-MØ and facilitates OxPhos, a metabolic pathway associated with IL-4-activated macrophages (Vats et al., 2006).

**Figure 2.**
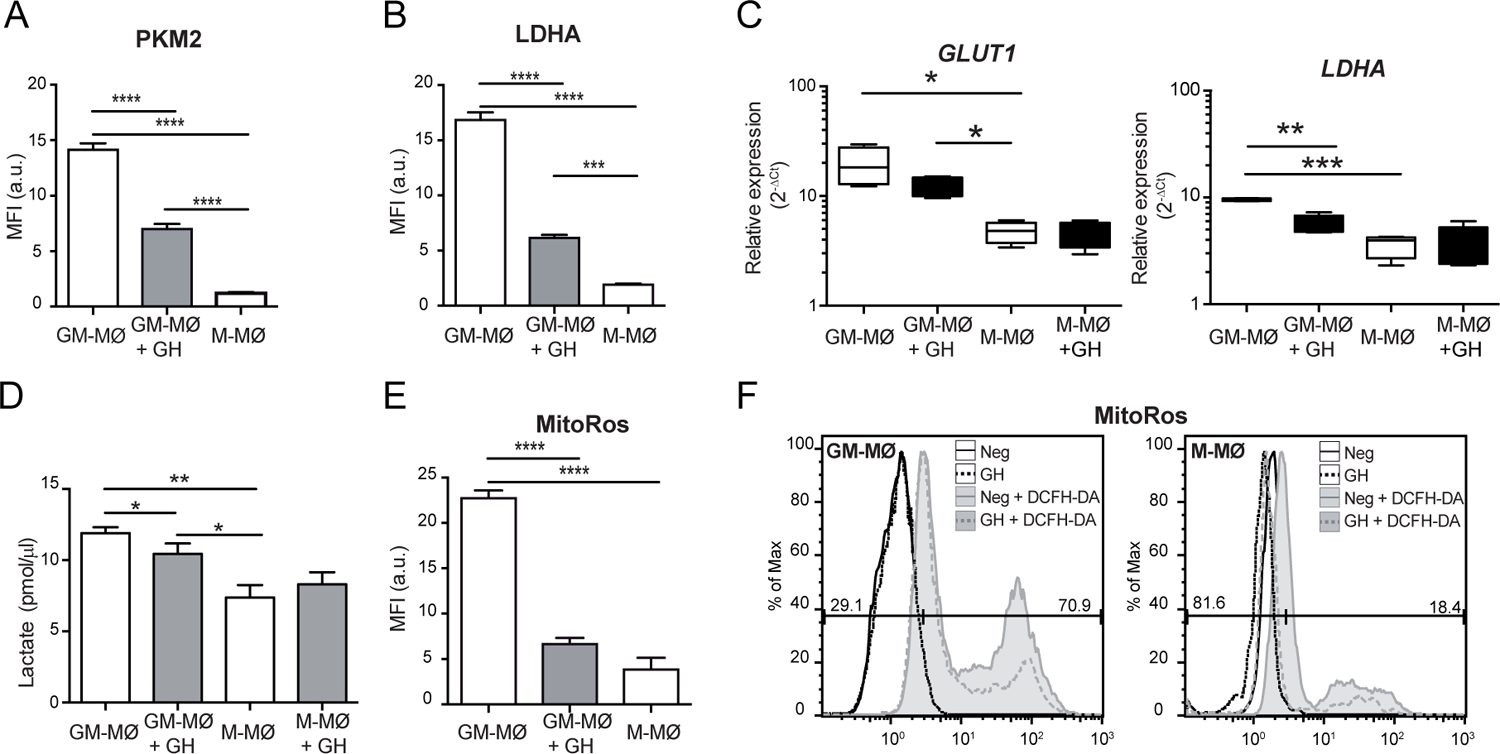
Treatment with rhGH decreases the expression of specific glycolysis enzymes and reduces lactate and ROS production in GM-MØ. Quantitative analysis of confocal images of GM-MØ treated or not with rhGH (1 μg/ml, 24 h) and M-MØ, stained with anti-PKM2 **(A)**, or -LDHA **(B)** monoclonal antibodies. Equatorial-plane images from 200–300 cells were collected in random fields and analyzed using ImageJ. Results are shown as mean ± SD after background substration (mean fluorescence intensity, MFI) (n=8). Paired *t*-test ***p<0.001, ****p<0.0001. **(C)** Relative expression levels of *GLUT1* and *LDHA* in GM-MØ and M-MØ treated or not with rhGH. Results are shown as 2^−ΔCt^ relative to the mean of internal *TBP* expression and correspond to triplicate determinations (n=8); box and whisker plots represent the median, second and third quartiles, and the minimum to maximum values. Paired *t*-test *p<0.05, **p<0.01, ***p<0.001. **(D)** Lactate determination in culture supernatants of GM-MØ and M-MØ treated or not with rhGH. Results are shown as mean ± SD (n=4). Paired *t*-test *p<0.05, **p<0.01. **(E)** Quantitative analysis of confocal images of GM-MØ treated or not with rhGH (1 μg/ml, 24 h) and M-MØ stained with MitoRos as in (A) (n=8). Results are shown as mean ± SD (n=4). Paired *t*-test ***p<0.001, ****p<0.0001. **(F)** ROS levels in GM-MØ and M-MØ treated or not with rhGH determined by incubation with DCFH-DA for 60 min followed by flow cytometry analysis. Representative plots are shown (n=8).

The TCA cycle plays an important role in cell metabolism and begins with the generation of citrate, which is formed by the condensation of oxaloacetate with acetyl-CoA generated from fatty acids, amino acids or pyruvate oxidation and processed in the cytosol by ACLY (ATP-citrate lyase). We thus examined the expression of ACLY and found lower expression in rhGH-treated GM-MØ (Fig. 3A, Supplemental Fig. 1A), which would suggest citrate accumulation in the cytosol. We also found that rhGH treatment significantly increased the gene expression levels of *CS* (citrate synthase) in GM-MØ but not in M-MØ (Fig. 3B, left), which would contribute to increase the mitochondrial levels of citrate. Citrate is converted to *cis*-aconitate, a metabolic intermediate of itaconate and succinate, by the action of aconitase. We observed that rhGH treatment reduced the expression of *IRG1* (aconitate decarboxylase 1), which is involved in itaconate production, in GM-MØ but not in M-MØ (Fig. 3Β, right), concomitant with a decrease in itaconate levels (Fig. 3C). These results agree with the increase in IDH2 (isocitrate dehydrogenase (NADP(+)) 2) and SDH (succinate dehydrogenase) enzyme levels by rhGH in GM-MØ (Fig. 3D; Supplemental Fig. 1A).

**Figure 3.**
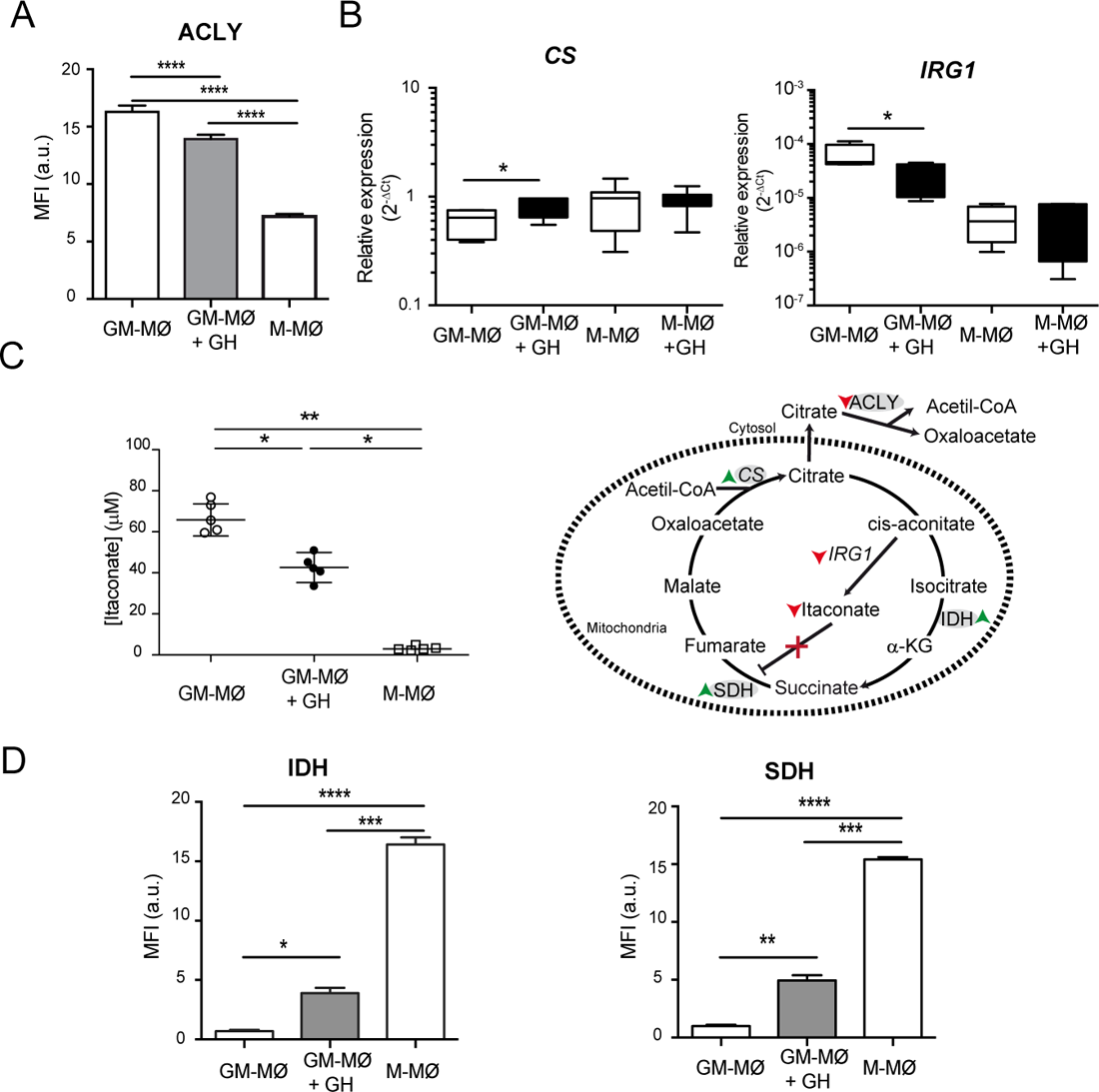
Modulation of itaconate levels in GM-MØ by rhGH. **(A)** Quantitative analysis of confocal images of GM-MØ treated or not with rhGH (1 μg/ml, 24 h) and M-MØ, stained with anti-ACLY monoclonal antibody. Equatorial-plane images from 200–300 cells were collected in random fields and analyzed using ImageJ. Results are shown as mean ± SD after background subtraction (mean fluorescence intensity, MFI) (n=8). Paired *t*-test ***p<0.001, ****p<0.0001. **(B)** *CS* and *IRG1* mRNA expression levels determined by RT-qPCR in untreated or rhGH-treated GM-MØ and M-MØ. Results are shown as 2^−ΔCt^ relative to the mean of internal *TBP* expression and correspond to triplicate determinations (n=8); box and whisker plots represent the median, second and third quartiles, and the minimum to maximum values. Paired *t-*test *p<0.05. **(C)** Intracellular levels (n=4 cultures) of itaconate in GM-MØ, rhGH-treated GM-MØ and M-MØ. All plots represent mean ± s.e.m., AU = arbitrary units based on mass spectrometry peak area. Paired *t-*test *p<0.05, **p<0.01. Effects of GH treatment on enzymes and products of the TCA cycle. Red arrows indicate downregulation, green arrows indicate upregulation and X represent blockade. **(D).** Quantitative analysis of confocal images of GM-MØ treated or not with rhGH (1 μg/ml, 24 h) and M-MØ stained with anti-IDH and -SDH monoclonal antibodies. Equatorial-plane images from 200–300 cells were collected in random fields and analyzed using ImageJ software. Results are shown as mean ± SD after background-subtraction. Paired *t*-test *p<0.05, **p<0.01, ***p<0.001, ****p<0.0001.

### Growth hormone affects the total mass of mitochondria, as well as their morphology and dynamics, in GM-MØ

In addition to their key role in cellular bioenergetics, mitochondria also produce metabolic precursors for macromolecules such as lipids, proteins and nucleic acids, and generate metabolic by-products such as ROS and ammonia (Spinelli & Haigis, 2018). As mitochondria are dynamic organelles that fuse and divide according to the metabolic and physiological needs of the cell, we hypothesized that GH might affect the total mass, dynamics and structure of mitochondria in GM-MØ. We thus stained untreated or rhGH-treated GM-MØ and M-MØ with MitoTracker Green (used to assess mitochondria mass) for confocal microscopy analysis. We found that the mass of mitochondria was significantly higher in GM-MØ than in M-MØ and rhGH treatment significantly decreased mitochondria mass in GM-MØ (Fig. 4A, Supplemental Fig. 1A).

**Figure 4.**
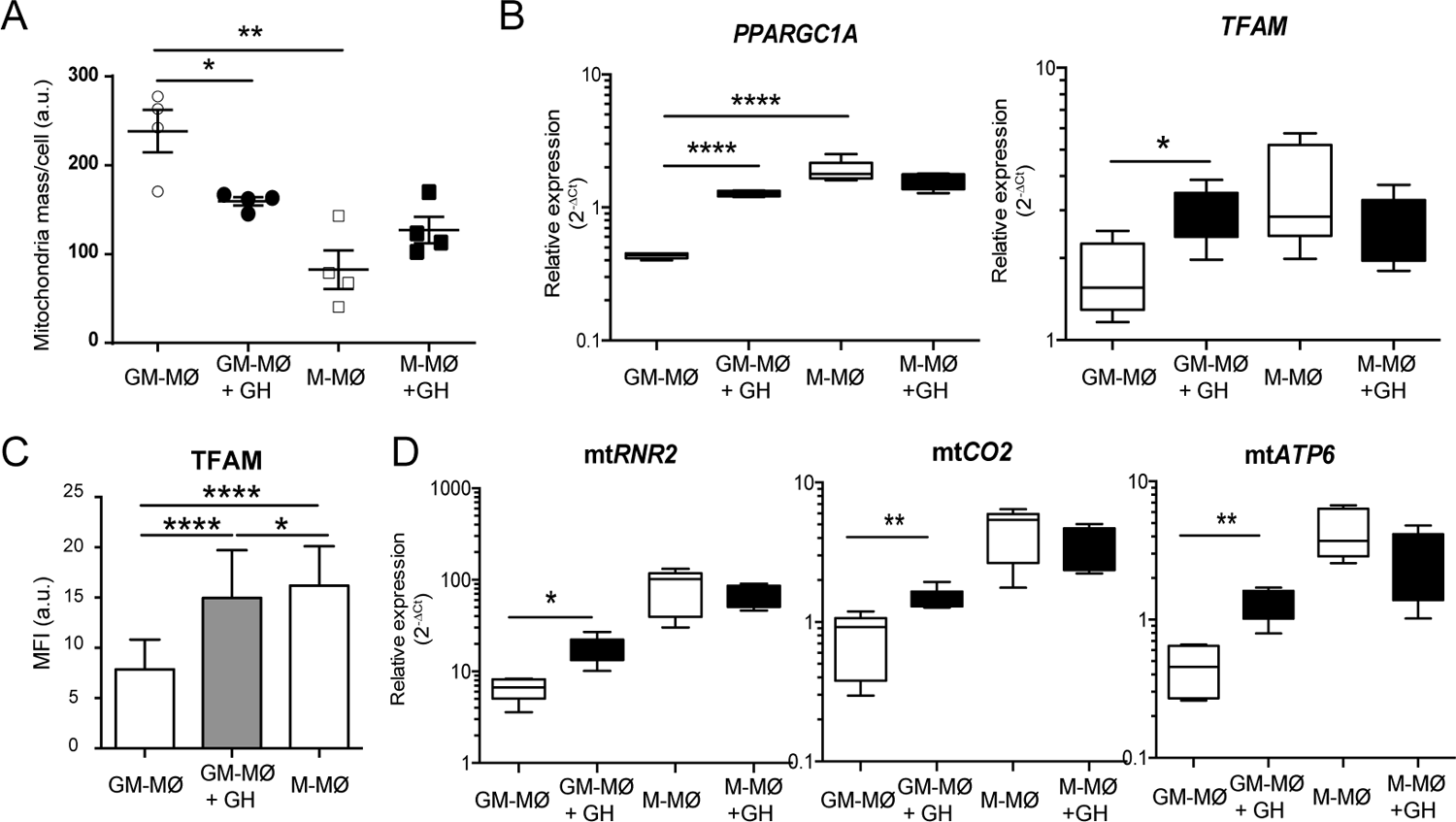
Treatment of GM-MØ with rhGH modulates mtDNA transcriptional activity. **(A)**. Quantitative analysis of confocal microscopy images of mitochondria using the MitoTracker Green probe. Data correspond to triplicate determinations (n = 8); box and whisker plots represent median, second and third quartiles, and the minimum to maximum values. Paired *t-*test *p<0.05, **p<0.01. **(B)**. *PPARGC1A* and *TFAM* mRNA expression levels determined by RT-qPCR in untreated or rhGH-treated GM-MØ and M-MØ. Results are shown as 2^−ΔCt^ relative to the mean of internal *TBP* expression and correspond to triplicate determinations (n=8); box and whisker plots represent the median, second and third quartiles, and the minimum to maximum values. Paired *t*-test *p<0.05, ****p< 0.0001. **(C).** Quantitative analysis of confocal images of GM-MØ treated or not with rhGH (1 μg/ml, 24 h) and M-MØ stained with anti-TFAM monoclonal antibody. Equatorial-plane images from 200–300 cells were collected in random fields and analyzed using ImageJ software. Results are shown as mean ± SD after background-subtraction. Paired *t*-test *p<0.05, ****p<0.0001. **(D)** *RNR2*, *CO2* and *ATP6* mitochondrial mRNA expression levels determined by RT-qPCR in untreated or rhGH-treated GM-MØ and M-MØ. Results are shown as 2^−ΔCt^ relative to the mean of internal *TBP* expression and correspond to triplicate determinations (n=8); box and whisker plots represent the median, second and third quartiles, and the minimum to maximum values. Paired *t*-test *p<0.05, **p<0.01.

Mitochondria abundance is regulated through biogenesis, fusion and fission events, and mitophagy. A key protein for mitochondrial biogenesis is peroxisome proliferator-activated receptor gamma coactivator 1α (PPARGC1α). Its activation triggers the subsequent activation of transcriptional regulators including nuclear respiratory factors (*NRF1* and *2*) and peroxisome proliferator-activated receptors (PPARs) (Scarpulla, 2008), which initiate transcription of nuclear genes involved in mitochondrial biogenesis and function. RT-qPCR analysis of untreated and rhGH-treated GM-MØ and M-MØ showed that *PPARGC1α* relative expression was significantly higher in M-MØ than in GM-MØ, and rhGH treatment significantly upregulated its expression in GM-MØ but not in M-MØ (Fig. 4B, left). A similar result was found for the transcription factor TFAM, which regulates mitochondrial biogenesis, both at the mRNA and protein level (Fig. 4B, right, C, Supplemental Fig. 1A). TFAM is abundantly expressed in human mitochondria, as it coats the entire mitochondrial DNA (mtDNA), and serves essential roles in mtDNA transcription (Maniura-Weber et al., 2004), replication (Pohjoismäki et al., 2009), maintenance (Kanki et al., 2004), packaging (Alam et al., 2003), and in nucleoid formation. Supporting this result, qRT-PCR analysis of untreated and rhGH-treated GM-MØ and M-MØ revealed a significant increase in the expression level of the mitochondrial genes *MT-CO2*, *MT-RNR2* and *MT-ATP6* in rhGH-treated GM-MØ but not in rhGH-treated M-MØ (Fig. 4D), confirming the effect of GH on the transcriptional activity of mtDNA in GM-MØ.

We next determined the effect of rhGH on mitochondria dynamics by evaluating fusion and fission processes. Untreated and rhGH-treated GM-MØ and untreated M-MØ (used as a control) were fixed and stained with a monoclonal antibody to the mitochondrial membrane protein TOM22 (Valente et al., 2017). In agreement with previous results in LPS/IFN-activated macrophages (Ramond et al., 2019), we observed that mitochondria (and mean branch length) were shorter in GM-MØ than in M-MØ (fission profile), which correlates with their glycolytic profile. Conversely, the elongated mitochondria in M-MØ (fusion profile) concurred with their higher OCR/ECAR ratio. Notably, rhGH treatment of GM-MØ decreased fission and increased fusion (Fig. 5A–C), in agreement with an effect of GH on reprogramming the metabolic profile of inflammatory macrophages (Nagdas & Kashatus, 2017). By contrast, rhGH failed to modify fission processes in M-MØ (Fig. 5A, B). The effect of rhGH on GM-MØ was also abrogated by rapamycin (Fig. 5A, C), confirming the involvement of mTOR pathway in the effect of rhGH on GM-MØ

**Figure 5.**
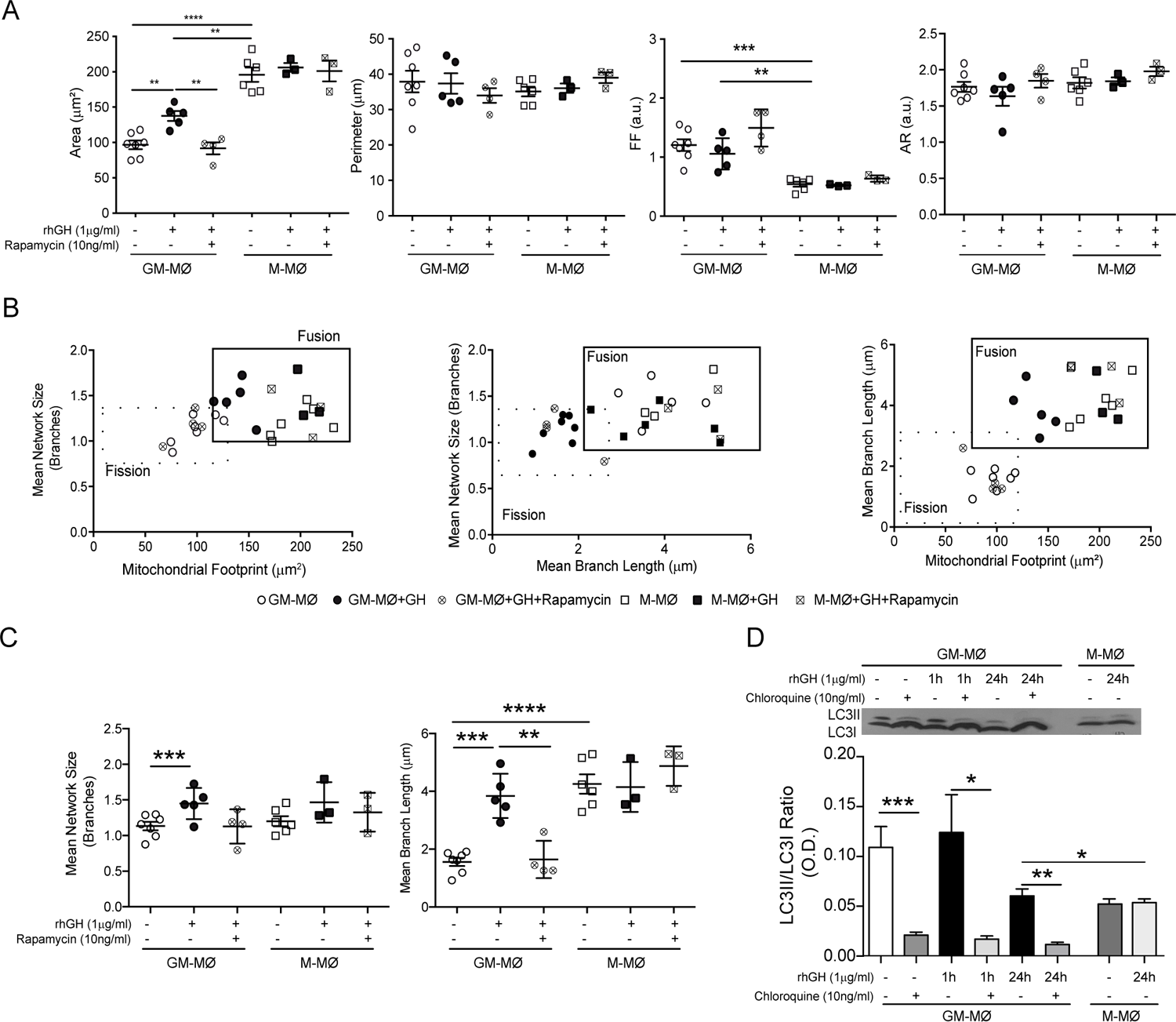
Treatment of GM-MØ with rhGH modulates mitochondrial dynamics. **(A)**. Quantitative analysis using specific macro tools of ImageJ of the area, perimeter, Form Factor (FF) and Aspect Ratio (AR) of untreated or rhGH-treated GM-MØ and M-MØ that were or not pretreated with rapamycin as indicated. The y-axis in each dot represents the feature values of individual cells (n≥8). Line and error bars represent mean and standard error of mean. Paired *t*-test *p<0.05, **p<0.01, ***p<0.001, ****p<0.0001. **(B).** Two-dimensional scatter plots from mitochondrial network analysis (footprints, network size, and branch length) matching data of individual cells plotted in (A). The quadrants of fission/fusion were added to enclose clustered macrophages in feature space based on their mitochondrial organization. Dots are data of individual cells (n≥8). All data shown were representative of three independent experiments. **(C).** Quantitative analysis of branch length of mitochondria from cells as in (A). Dots are data from individual cells (n≥8) of three independent experiments performed. Results are mean ± SD. Paired *t*-test **p<0.01, ***p<0.001, ****p<0.0001. **(D).** Representative western blot analysis of LC-3II and LC-3I expression in lysates of untreated or rhGH-treated GM-Mϕ and M-Mϕ at different time points. As control the effect of chloroquine (10 ng/ml, 30 min) pretreatment on untreated or rhGH-treated GM-Mϕ was also shown. Densitometric analysis of LC-3II and LC-3I levels normalized to those of vinculin. Mean ± SD (n=4). Paired *t*-test **p<0.01, ****p<0.0001.

In addition to mitochondrial biogenesis and dynamics, mitophagy is also essential to maintain overall mitochondrial homeostasis by selectively removing aged and damaged mitochondria *via* the specific sequestration and engulfment of mitochondria for subsequent lysosomal degradation in a process dependent on LC3-mediated autophagosome formation. Western blot analysis of GM-MØ and M-MØ showed a higher abundance of LC3-II-positive autophagosomes in inflammatory (GM-MØ) macrophages than in M-MØ. We found that rhGH treatment of GM-MØ suppressed LC3-II accumulation in a time-dependent manner (Fig. 5D). As a control for this analysis we used chloroquine, a drug that inhibits autophagic degradation in lysosomes (Mizushima & Yoshimori, 2007), which reduced LC3-II accumulation in GM-MØ. These data suggest that mitophagy events are more common in inflammatory than in anti-inflammatory macrophages and that rhGH treatment of GM-MØ likely abrogates this process. Overall, our findings indicate that GH might regulate the efficient production of ATP and the production of lipids and proteins in GM-MØ by affecting both the mass and dynamics of mitochondria.

Finally, we combined confocal live-cell imaging with correlative cryogenic fluorescence microscopy and cryo-FIB-SEM volume imaging to analyze the effect of rhGH on the shape and 3D-organization of mitochondria and their cristae in GM-MØ. GM-MØ, M-MØ and rhGH-treated GM-MØ were seeded on finder gold grids and then stained with MitoTracker Red FM (a mitochondrial membrane-dependent dye) prior to sample vitrification and evaluation by cryo-fluorescence microscopy. Analysis indicated that mitochondrial volume was smaller in GM-MØ than in M-MØ and that rhGH-treatment of GM-MØ significantly increased mitochondrial volume (Fig. 6A). Additionally, we confirmed that GM-MØ mitochondria harbored a fission profile, whereas M-MØ mitochondria harbored a fusion profile, and that rhGH treatment of GM-MØ reduced fission and increased fusion events (Supplementary Fig. 4A, B), an effect that was also evident when the elongation coefficient of the mitochondria was determined. Indeed, elongation was significantly higher in M-MØ and in rhGH-treated GM-MØ than in untreated GM-MØ (Supplementary Fig. 4C). We also performed a quantitative analysis of the mitochondrial ultrastructure by determining the ratio between the cristae volume and the mitochondria volume as a measurement of the density and/or volume of cristae per mitochondria. The results indicated a higher density of cristae in anti-inflammatory (M-MØ) macrophages than in inflammatory macrophages (GM-MØ), and that rhGH-treatment increased the density of cristae in GM-MØ (Fig. 6B, C). These data correlate well with the upregulation of OxPhos activity and with the increased mtDNA copy number and transcription levels of mtDNA in rhGH-treated GM-MØ.

**Figure 6.**
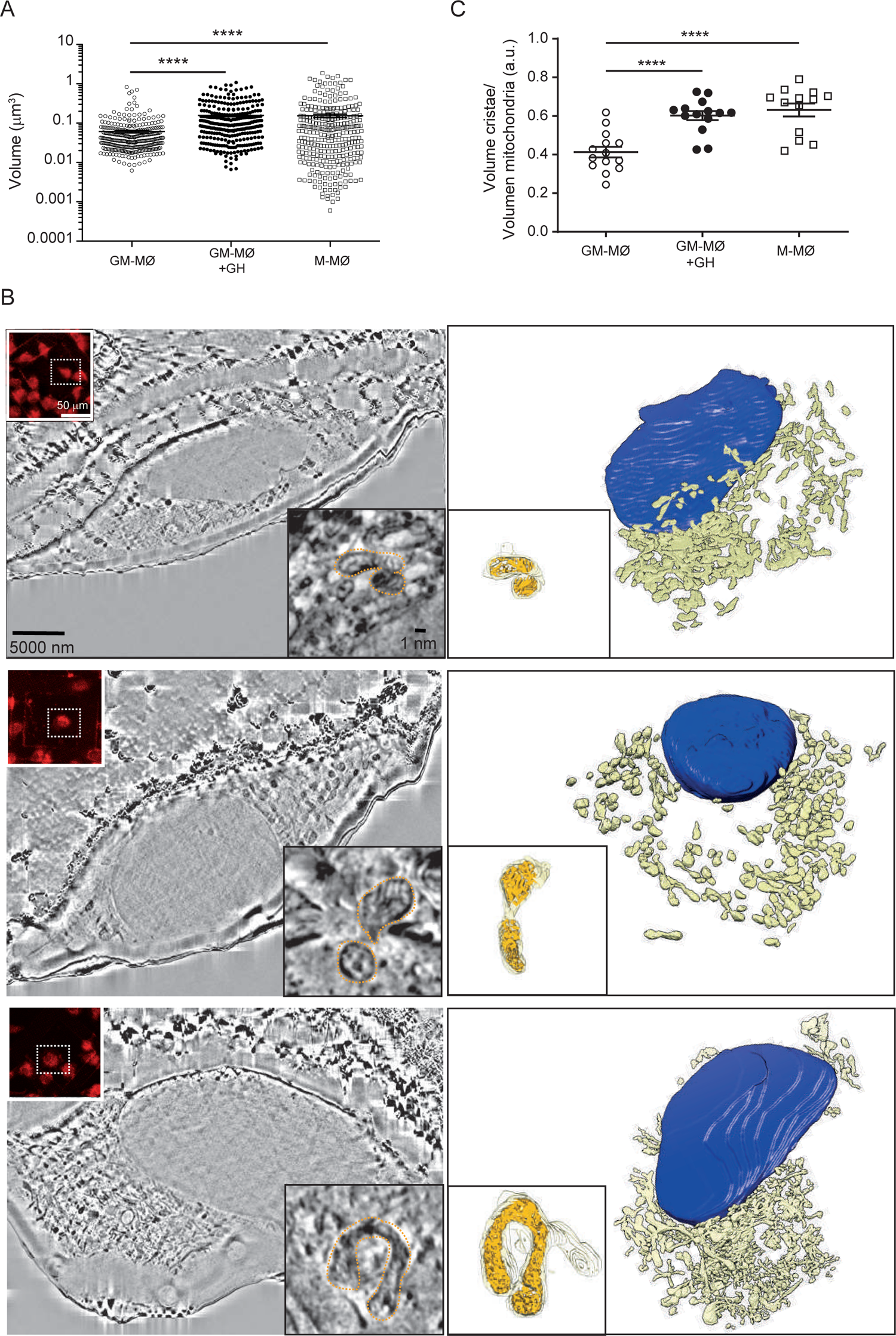
Treatment with rhGH alters GM-Mϕ mitochondria intimal structure. **(A)** Mitochondrial volume from each experimental condition, untreated or rhGH-treated GM-MØ and M-MØ. Line and error bars represent mean and SEM. Paired *t* test ****p<0.0001. **(B)** Reconstruction of untreated GM-MØ (upper), rhGH-treated GM-MØ (middle) and untreated M-MØ (lower). In all cases, a representative cryo-FIB-SEM 3D section is shown (left) with a confocal microscopy image of the MitoTracker signal (treated 30 min at 37°C) (upper left corner inset), a detailed image of one of the layers of the mitochondrion used to quantitate the volumes (right lower corner inset). Figure also shows the corresponding 3D segmentation (right) with a 3D representation of a mitochondrion (lower left corner inset) surface (yellow) and its inner structure (orange). **(C)**. Quantitation of the mitochondrial cristae volume relative to the total volume of the mitochondria corresponding to untreated GM-MØ, rhGH-treated GM-MØ and untreated M-MØ. Three representative mitochondria from each treatment per cell were analyzed (4 cells per donor, 2 different donors). Line and error bars represent mean and standard error of mean. Paired *t-*test ns not significant ****p<0.0001.

## Discussion

Macrophages comprise a functionally heterogeneous cell population with vital roles in host defense against pathogenic infections and in inflammatory responses. Their extreme phenotypic plasticity allows them to adapt their responses to multiple microenvironmental cues. Classically-activated (M1) macrophages are potent effector cells that kill microorganisms and secrete inflammatory factors, whereas alternatively-activated (M2) macrophages scavenge debris, promote allergy reactions, participate in tissue remodeling and repair, and secrete anti-inflammatory cytokines (Wynn et al., 2013). Both types of macrophages use distinct pathways for energy production: whereas LPS/IFN-activated macrophages have enhanced glucose consumption and lactate release, IL4-activated macrophages mainly use oxidative glucose metabolism (Odegaard & Chawla, 2011; Rodríguez-Prados et al., 2010). Metabolic requirements influence macrophage plasticity, and their metabolic reprogramming regulates how they execute specific effector functions. Thus, instructing macrophage metabolism is a potential strategy to modulate their activity, which might be useful in diseases with a high macrophage commitment (O’Neill et al., 2016; O’Neill & Pearce, 2016). For example, by promoting the polarization of inflammatory macrophages in atheroma plaques towards an anti-inflammatory phenotype, IL-13 reduces atherosclerosis (Stöger et al., 2012). Similarly, decreasing lipid levels or enhancing HDL levels switches inflammatory macrophages to an anti-inflammatory phenotype in atheroma plaques and triggers atherosclerosis regression (Feig et al., 2011).

While GM-CSF- and M-CSF-derived macrophages require LPS/IFN or IL-4 activation, respectively, to generate classically- or alternatively-activated macrophages, analysis of their gene expression profile indicates that GM-CSF and M-CSF prime macrophages towards M1 and M2 phenotypes and, therefore, they can be considered inflammatory or anti-inflammatory macrophages (Lacey et al., 2012; Martinez et al., 2006). We recently reported that GH reprograms GM-MØ toward an anti-inflammatory and reparative profile (Soler Palacios et al., 2020). Supporting this, rhGH treatment of GM-MØ promoted a significant enrichment of anti-inflammatory gene expression and dampened the proinflammatory cytokine profile, which was in agreement with the improved remission of inflammation and mucosal repair during recovery in the acute dextran sodium sulfate–induced colitis model in bovine GH (bGH)-overexpressing mice (Soler Palacios et al., 2020). We hypothesized that rhGH treatment of human GM-MØ reprograms their metabolism, relatively suppressing glycolysis and lactate accumulation in favor of OxPhos. Whole cell respiration data support this hypothesis and protein expression analysis revealed a rhGH-mediated downregulation of PKM2 and LDHA, two key proteins involved in glycolysis. By contrast, we failed to observe any effect of rhGH on M-MØ metabolism. Confirming its role in suppressing glycolysis in GM-MØ, rhGH treatment reduced the mRNA levels of the glucose transporter *GLUT1*. GLUT1 is the primary rate-limiting glucose transporter in proinflammatory-polarized macrophages and its rapid upregulation is associated with a proinflammatory phenotype (Freemerman et al., 2014). This observation correlates with published data showing that hGH reduces the synthesis of GLUT1 and inhibits 2-deoxyglucose and 3-O-methyl-D-glucose uptake in 3T3-F442A adipose cells (Ku Tai et al., 1990).

Consistent with the decrease in LDHA expression by rhGH, we also detected a decline in lactate accumulation in GM-MØ. Lactate has a critical function in inducing M2-like polarization in tumor models (Colegio et al., 2014). In LPS/IFN-activated macrophages, the generation of lactate *via* pyruvate is essential to restore NAD^+^ levels and maintain flux through the glycolytic pathway (Viola et al., 2019). We speculate that the observed drop in lactate levels after rhGH treatment of GM-MØ correlates with the hormone-triggered downregulation of glycolysis and ROS production. These observations accord with rhGH favoring the OxPhos pathway in inflammatory macrophages. We also detected that rhGH treatment decreased ACLY levels in GM-MØ. ACLY has been associated with activated inflammatory macrophages and is related to the presence of human atherosclerotic plaques. ACLY deficiency in myeloid cells induces a stable plaque phenotype, as demonstrated by increased collagen content and fibrous cap thickness, along with a smaller necrotic core size (Baardman et al., 2020). In humans, severe GH deficiency is associated with increased cardiovascular risk and intima-media thickness at major arteries, a phenotype that can be reversed with rhGH replacement therapy (Colao et al., 2008). We also detected lower levels of intracellular itaconate and of *IRG1*, which encodes the enzyme that converts *cis*-aconitate to itaconate, in rhGH-treated inflammatory macrophages. Itaconate accumulation is a prime indicator of metabolic reprogramming in macrophages upon LPS treatment (Lee et al., 1995) and it is consistently detected in inflammatory macrophages where it is known to play an immunomodulatory role (Hooftman et al., 2020), promoting anti-inflammatory functions (M. P. Murphy & O’Neill, 2018).

Mitochondria are signaling organelles that regulate a wide variety of cellular functions and can dictate cell fate. Their structure is highly dynamic and their organization can determine the metabolic function of cells (Archer, 2013). Several reports have produced controversial observations relating mitochondrial mass to inflammation. For example, it has been shown that an increase in mitochondrial mass is necessary for pro-inflammatory differentiation of macrophages (Yu et al., 2020), and that mitochondrial mass may be critical for the control of cell fate and immune responses. Also, higher mitochondrial mass has been shown to contribute to T-cell senescence and cancer cell chemo-resistance (Callender et al., 2020; Davizon-Castillo et al., 2019), whereas PGC-1α-deficient mice show a lower mitochondrial mass, which causes spontaneous kidney inflammation and injury (Fontecha-Barriuso et al., 2019), and LPS treatment increases mitochondrial mass in macrophages together with pro-inflammatory cytokine production (Yu et al., 2020). We found that mitochondria mass was significantly lower in rhGH-treated GM-MØ than in equivalent untreated cells. Mitochondria continuously remodel their structure through biogenesis and mitophagy, and alternate processes of fission and fusion. Our results indicate that rhGH treatment stimulates mitochondria biogenesis, reduces fission and increases fusion events and reduces mitophagy in inflammatory GM-MØ. Fusion events are associated with increased ATP production, activated OxPhos and ROS production (Wai & Langer, 2016). Mitochondrial fusion is also required to maintain the stoichiometry of the mtDNA replisome and the integrity of the mitochondrial genome (Chapman et al., 2020). We also found an upregulation of TFAM in rhGH-treated GM-MØ. TFAM is a multifunctional DNA-binding protein that is essential for transcriptional activation and mtDNA organization (Bonawitz et al., 2006). Mice lacking TFAM have impaired mtDNA transcription and are unable to maintain mtDNA, resulting in bioenergetic failure and embryonic lethality (Larsson et al., 1998). rhGH-mediated TFAM upregulation correlated with the increased expression levels of the mitochondrial genes *CO2*, *RNR2* and *MT*-*ATP6*.

Analysis of mitochondria shape and internal structure, using correlative cryogenic fluorescence microscopy combined with cryo FIB-SEM, indicated a rhGH-mediated increase of GM-MØ mitochondria volume and elongation, which concurred with the increase in fusion events. We also detected higher density of mitochondrial cristae in GM-MØ treated with rhGH, in agreement with the metabolic shift towards the preferential OxPhos activity and ATP production detected in anti-inflammatory macrophages.

mTOR signaling is the most important intracellular pathway coordinating cellular metabolism (Saxton & Sabatini, 2017), as it stimulates glycolysis and glucose uptake through the transcription factor hypoxia-inducible factor-1 alpha (Düvel et al., 2010). In particular, the PI3K/Akt/mTOR signaling pathway is essential for macrophage activation, controlling either canonical signaling (e.g., JNK, NF-kB) or metabolic processes (Covarrubias et al., 2015). We observed that the rhGH-mediated effects on mitochondria dynamics were abrogated in rapamycin-treated GM-MØ, in agreement with our previous results showing that rhGH-mediated reprogramming of GM-MØ is a pAKT-mediated process (Soler Palacios et al., 2020) and with reports demonstrating that mTORC1 (mTOR complex1) is downstream of AKT (Kim & Guan, 2019).

Overall, our results support a modulatory function of GH in the metabolism of the inflammatory macrophages that might contribute to immune regulation directly or indirectly through metabolic coupling between macrophages and other cell types. As macrophage metabolism is inextricably linked to their functionality, metabolic reprogramming of macrophages might be an elegant way to intervene in multiple inflammatory diseases such as atherosclerosis or adipose tissue obesity-induced insulin resistance (Geeraerts et al., 2017). Epidemiological evidence suggests that aging is the single biggest risk factor for chronic inflammatory diseases and, mechanistically, inflammation is thought to be a common link between aging and disease. Our results thus suggest that GH treatment may also contribute to restore immune system homeostasis and control age-related inflammation. The metabolism of macrophages, and, in particular their metabolic repolarization, thus offers new therapeutic opportunities to treat inflammatory diseases and cancer.

## Acknowledgments

We thank Dr. A.L. Corbí from the Myeloid Cell Biology Group, Centro De Investigaciones Biológicas Margarita Salas/CSIC Madrid, Spain for critical review of the manuscript. We are grateful to Pfizer for the gift of Genotonorm® (CPT grant WI215392). We also acknowledge access to the cryoEM CNB-CSIC facility in the context of the CRIOMECORR project (ESFRI-2019-01-CSIC-16). This work was supported in part by grants from the Spanish Ministry of Science and Innovation (PID2020-114980RB-I00) Agencia Estatal de Investigación/Fondo Europeo de Desarrollo Regional (AEI/FEDER), Unión Europea. AC was supported by an FPI fellowship from the Spanish Ministry of Science and Innovation (PRE2018-086112) through the Severo Ochoa excellence accreditation SEV-2017-0712-18-1.

## Material and Methods

### Cell culture

Human peripheral blood mononuclear cells were isolated from buffy coats from healthy donors over a Lymphoprep (Nycomed Pharma AS, Oslo, Norway) gradient according to standard procedures, and monocytes were then purified using magnetic cell sorting with CD14 microbeads (Miltenyi Biotech GmbH, Bergisch Gladbach, Germany). Monocytes were cultured at 0.5 × 10^6^ cells/ml for 7 days at 37°C in a humidified atmosphere with 5% CO_2_ in RPMI medium containing 10% fetal calf serum, and supplemented with GM-CSF (1000 U/ml) or M-CSF (10 ng/ml) (both from ImmunoTools GmbH, Friesoythe, Germany) to generate GM-CSF-polarized macrophages (GM-MØ) or M-CSF-polarized macrophages (M-MØ). rhGH (Pfizer Genotonorm®, 1 µM) was added to 7-day differentiated macrophages for 24 h. When required, 5 × 10^6^ cells were treated with chloroquine (10 ng/ml, 30 min, 37°C; Sigma-Aldrich, Madrid, Spain) and 1 × 10^6^ cells were treated with rapamycin (10 ng/ml, 30 min, 37°C; Calbiochem, San Diego, CA) prior to treatment with rhGH.

### Gene-set enrichment analysis

RNA-sequencing data have been deposited in the Sequence Read Archive (SRA) (https://www.ncbi.nlm.nih.gov/sra) under accession no. PRJNA555143. GSEA was performed using the data sets Hallmark Glycolysis; http://www.gsea-msigdb.org/gsea/msigdb/cards/HALLMARK_GLYCOLYSIS.html (Liberzon et al., 2015), available at http://software.broadinstitute.org/gsea/index.jsp.

### Measurement of cellular respiration and extracellular acidification

The bioenergetic profile of cells was measured by determining the OCR and the ECAR on the Seahorse XF-24 extracellular flux analyzer platform (Seahorse Biosciences, North Billerica, MA). Cells were seeded (2 × 10^5^ cells/well) in XF-24 plates and allowed to recover (24 h, 37°C). Cells were then incubated in bicarbonate-free DMEM (Sigma-Aldrich) supplemented with 11.11 mM glucose, 2 mM L-glutamine, 1 mM pyruvate, and 2% fetal bovine serum (Sigma-Aldrich) in a CO_2_-free incubator (1 h, 37°C). After four measurements under basal conditions, cells were treated sequentially with 1 mM oligomycin, 0.6 mM carbonyl cyanide p-(trifluoromethoxy) phenylhydrazone (FCCP), 0.4 mM FCCP, and 0.5 mM rotenone plus 0.5 mM antimycin A (Sigma-Aldrich), with three consecutive determinations under each condition, which were subsequently averaged. Non-mitochondrial respiration (OCR value after rotenone plus antimycin A addition) was subtracted from all OCR measurements. Basal OCR corresponds to the OCR in the absence of effectors. ATP turnover was estimated from the difference between the basal and the oligomycin-inhibited respiration, and the maximal respiratory capacity was the rate in the presence of the oxidative phosphorylation uncoupler FCCP. Eight independent biological replicates of each analysis were performed, and results were normalized to protein concentrations as determined by BCA assays.

### Immunohistochemistry and multicolor confocal microscopy

The following antibodies were used: anti-PKM2 (sc365684; Santa Cruz Biotechnology, Santa Cruz, CA), -LDHA (2012; Cell Signaling Technology, Danvers, MA), -ACLY (ab407793; Abcam, Cambridge, UK), -SDH (CL0349; ThermoFisher Scientific, Waltham, MA), -IDH (8H26L4; ThermoFisher Scientific), -TFAM (16832595; ThermoFisher Scientific), -H3K4me3 (ab8580; Abcam) and isotype-matched control antibodies and fluorochrome-conjugated secondary antibodies (Jackson ImmunoResearch Laboratories, West Grove, PA). Macrophages were plated on poly-L-lysine (Sigma-Aldrich)-coated coverslips, fixed with 4% formaldehyde and, when indicated, permeabilized with 0.1% saponin for 10 min. Cells were then blocked for 10 min with 1% human immunoglobulins before incubation with primary (1–5 μg/ml) and appropriate secondary antibodies.

For *in vivo* experiments, macrophages were incubated in the complete RPMI containing 100 nM MitoTracker FITC (11589106; Invitrogen, Carlsbad, CA), MitoROS-FITC (25169; Cayman Chemical, Ann Arbor, MI) and CMXRos-Red (M7512; ThermoFisher Scientific) for 30 min and then with 2 µg/mL DAPI (Sigma-Aldrich) for an additional 2 min. After incubation, cells were washed twice and resuspended in complete RPMI. For live-cell imaging, macrophages seeded on confocal dishes were placed in a micro-incubator system of 37°C and 5% CO_2_ in a humidified environment. Imaging was performed with an inverted confocal microscope (SP5; Leica Microsystems, Buffalo Grove, IL), using the 20 PL-APO NA 0.7 and the 63 PL-APO NA 1.3 glycerol immersion objectives.

Quantification of protein expression was performed using similar acquisition settings in all cells and localized with DAPI and differential interference contrast. Mean fluorescence intensities (arbitrary units) within regions of interest (ROIs) were assessed using ImageJ (NIH, Bethesda, MD) (Schindelin et al., 2012). At least six independent samples were evaluated for each type of macrophage. After background subtraction, data were plotted using Prism software (GraphPad Software, La Jolla, CA,).

### Lactate quantitation

Supernatants were analyzed for the presence of lactate using the Lactate Colorimetric Assay Kit II (Quimigen, Madrid, Spain Catalog #K627-100).

### Detection of intracellular ROS

Cells were incubated with 20 µM H_2_DCF-DA (ab113851; Abcam) in culture medium for 30 min at 37°C and 10% CO_2_. The H_2_DCF-DA-loaded cells were washed, resuspended in RPMI and further incubated for 30 min. Cell fluorescence were quantified using a flow cytometry (FACS Aria, BD Biosciences, San Jose, CA,).

### Determination of intracellular itaconate levels

Intracellular itaconic acid levels were measured using high performance liquid chromatography. Pellets from 2 × 10^7^ macrophages were resuspended in PBS, pH 7.4 and lysed by three freeze-thaw cycles. Samples were bath-sonicated (5 min, room temperature) in two steps with an intermediate freeze step. After centrifugation (17, 000 × g, 15 minutes, 4°C), debris-free lysates were transferred to new tubes. Proteins were then precipitated using 100 mM hydrochloric acid and centrifuged (17, 000 × g, 15 min, 4°C). Remaining supernatants were injected into a column of 250 × 2 mm (Col Kromophase 100 C18 5.0 mm; Scharlau Chemie SA, Barcelona, Spain) and itaconate was quantified using UV detection at 210 mm. A standard curve (Supplemental Fig. 1C) was obtained using (0.50 μM to 500 μM) pure itaconic acid (Sigma-Aldrich, I29204). Results were normalized using the protein concentration of each sample.

### Quantitative real-time PCR

Total RNA from cultured cells was extracted using the Nucleospin RNA/Protein kit (Macherey-Nagel, Düren, Germany). RNA was retrotranscribed using SuperScript III reverse transcriptase (Invitrogen) and RT-qPCR analysis was performed with GoTaq qPCR Master Mix (Promega, Madison, WI). Triplicate samples were quantified using the QuantStudio 5 software (Applied Biosystems). Oligonucleotides for selected genes were designed employing the Roche Universal ProbeLibrary Assay Design Center and (Kraja et al., 2019) (Table 1). Assays were performed in triplicate and results were normalized according to the average expression levels of the housekeeping gene *TBP*. Results were obtained using the 2^-ΔCt^ comparative threshold method for quantification.

### Fusion-fission experiments and image analysis

Macrophages were plated on poly-L-lysine (Sigma-Aldrich)-coated coverslips, fixed with 4% formaldehyde and, when indicated, permeabilized with 0.1% saponin for 10 min. Cells were then blocked for 10 min with 1% human immunoglobulins, and incubated with TOM22 (1 µg/ml, Sigma-Merck; HPA003037) and appropriate secondary antibodies. Imaging was acquired with an inverted confocal microscope (YODA WF/TIRFM, Leica Microsystems), using the 20 PL-APO NA 0.7 and the 63 PL-APO NA 1.3 glycerol immersion objectives and an EM-CCD camera (Andor DU 885-CS0-#10-VP; Andor Technology, Belfast, UK).

The approach to quantitatively evaluate the mitochondrial characteristics is summarized in Supplemental Fig. 2. Analysis of mitochondrial morphology was performed using the macro complement Mitochondrial Flow of ImageJ (https://github.com/QuantitativeImageAnalysisUnitCNB/MitochondrialFlow_) (Valente et al., 2017). Briefly, images were first cropped to show a single cell per image. Next, selected cells were processed using an enhanced local contrast (CLAHE macro) and a binary image was obtained and processed using as threshold (Size = 0.06 µm^2^-Infinity, Circularity = 0.00–1.00), to finally obtain the Area, Perimeter, and Shape Descriptors values. The Form Factor (FF) data was calculated as the inverse of the “Circularity” output value. The network connectivity analysis was performed using the Skeletonize 2D to generate a skeleton map, and the Analyze Skeleton plugin to calculate the number of branches, branch lengths, and branch junctions.

### Western blotting

Cell lysates were obtained in RIPA buffer containing 2 mM Pefabloc, 2 mg/ml aprotinin/antipain/leupeptin/pepstatin, 10 mM NaF, and 1 mM Na_3_VO_4_. Cell lysates (30 μg/ml) was subjected to SDS-PAGE and transferred onto Immobilon polyvinylidene difluoride membranes (Millipore, Billerica, MA). Protein detection was carried out using anti-human LC3-I/II antibody (Cell Signaling Technology, Danvers, MA, #12741).

### Correlative cryogenic fluorescence microscopy and cryo FIB-SEM Vitrification

Cells (0.5 × 10^6^) were seeded on Quantifoil silicon oxide grids R 1/4 finder F1 Au grids, 200 mesh (Quantifoil Micro Tools GmbH, Jena, Germany). Cells were then cultured in RPMI (24 h, 37°C, 5% CO_2_) and stained with MitoTracker Red FM (ThermoFisher Scientific, M22425; 30 min, 37°C). Grids were vitrified by plunge-freezing using a Leica EM GP2 grid plunger (Leica Microsystems, Vienna, Austria) set to 37°C, 95% humidity and a blotting time of 7 s by the grid side opposite to the growing cells. Immediately before vitrification, 3 µl of Dynabeads MyOne Carboxylic Acid (1 μm; ThermoFisher Scientific) were added at a concentration of 0.5 mg/ml to each of the samples. Vitrified grids were mounted under liquid nitrogen in c-clip rings (ThermoFisher Scientific).

### Image acquisition

#### Cryo-fluorescence microscopy

The approach to quantitate the mitochondrial characteristics is summarized in Supplemental Fig. 3. Vitrified samples were analyzed by cryo-fluorescence microscopy using a LSM 900 confocal microscope (Carl Zeiss NTS GmbH, Oberkochen, Germany) equipped with a cryo-stage Linkam CSM196 (Linkam, UK). Fluorescence images were acquired with a LD A-Plan 20×/0.35 Ph1 objective using two channels, brightfield and red fluorescence emission, both with a pinhole aperture of 67 μm. Brightfield images were collected using an ESID detector (photodiode) with an excitation light of 400 nm. Red fluorescence images were collected using a GaAsP-PMT detector and an ex/em wavelength of 578/598 nm. Whole grids were imaged to generate a map of stitched images (9 images per grid) to screen grids and locate cellular coordinates of interest.

#### Cryo-FIB-SEM volume imaging

Cryo-fluorescence microscopy samples were transferred to the cryo-FIB-SEM microscope in liquid nitrogen using a Leica EM-VCM500 machine and a cryo-holder suitable for c-clipped grids (Leica Microsystems, Vienna, Austria). Prior to sample loading into the FIB-SEM microscope, grids were metalized by platinum sputtering (4 nm) using a Leica ACE600 cryo-sputter coater equipped with a cryo-stage (Leica Microsystems). Metallized samples were then transferred to a pre-cooled Crossbeam 550 FIB-SEM microscope (Carl Zeiss NTS GmbH) equipped with a cryo-stage (Leica Microsystems) using an EM-VCT100 shuttle (Leica Microsystems). The cryo-stage was positioned at a working distance of 5.1 mm and tilted to 11° and further protected by a layer of cold-deposited organic-platinum (3 depositions in cycles of 30 s not heating the source at a working distance of 8 mm). Serial sectioning of the sample was followed to generate the FIB-SEM volumes using SmartFIB software (Carl Zeiss NTS GmbH). SEM imaging conditions were 1.8 kV acceleration voltage and a probe current of 36 pA using the InLens detector and at a final pixel size between 4 and 9 nm. FIB milling was done at 30 kV and 300 pA with a sectioning step between 20 and 25 nm. Four different cells from each condition were imaged.

#### Image processing and segmentation

Raw FIB-SEM image stacks pre-processing was done using ImageJ. To remove the curtain artefact in the raw images a vertical band-pass filter between 1 and 300 pixels was applied. Images stack alignment was done using Linear Stack Alignment with SIFT (Lowe, 2004). Aligned stacks were then submitted to semi-manual segmentation and statistical analysis using Amira software (ThermoFisher Scientific). We obtained the morphological data of the volume and elongation of each mitochondrion in untreated or rhGH-treated macrophages. Elongation was defined as the ratio of the medium and the largest eigenvalue of the covariance matrix calculated for each mitochondrion. To evaluate fusion-fission events in 3D we used ImageJ. The approach for mitochondrial characteristics quantitation and subsequent morphology analysis was obtained using a public Java-based “MitochondrialAnalyzer” plugin operational under ImageJ software (https://github.com/QuantitativeImageAnalysisUnitCNB/MitochondrialAnalyzer_).

#### Statistical analysis

For comparison of mean or median, and unless otherwise indicated, statistical significance of the data was evaluated using a paired *t*-test. A p-value <0.05 was considered significant (*p<0.05, **p<0.01, ***p<0.001, ****p<0.0001). Statistical parameters used in the GSEA analysis were as previously described (Subramanian et al., 2005).

---

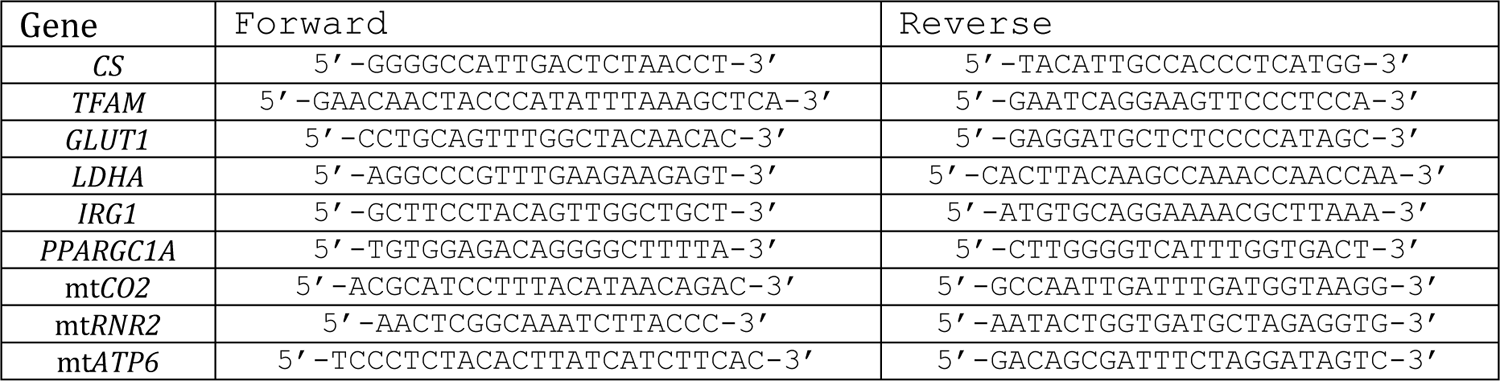

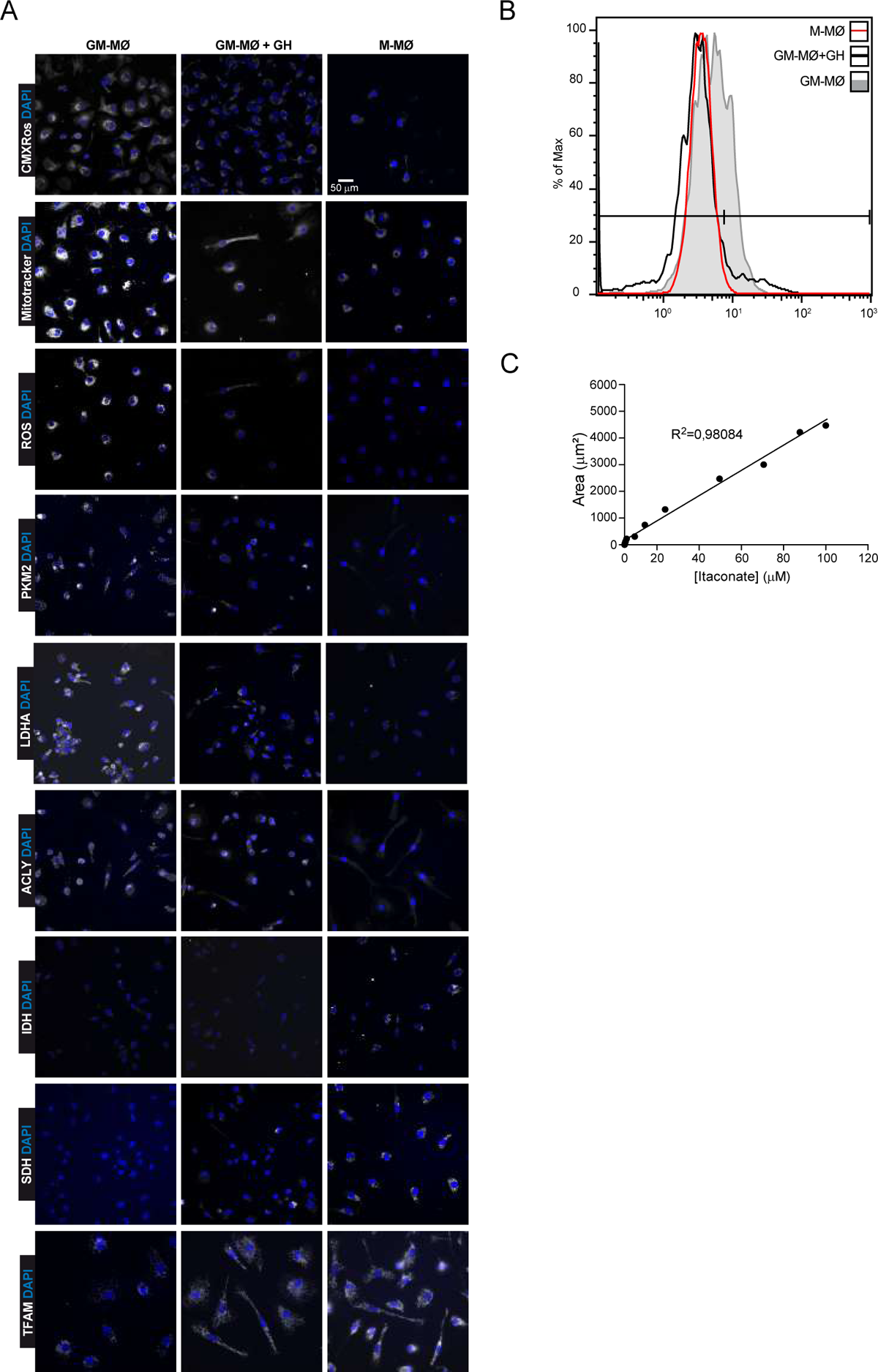

**Supplementary 2.**
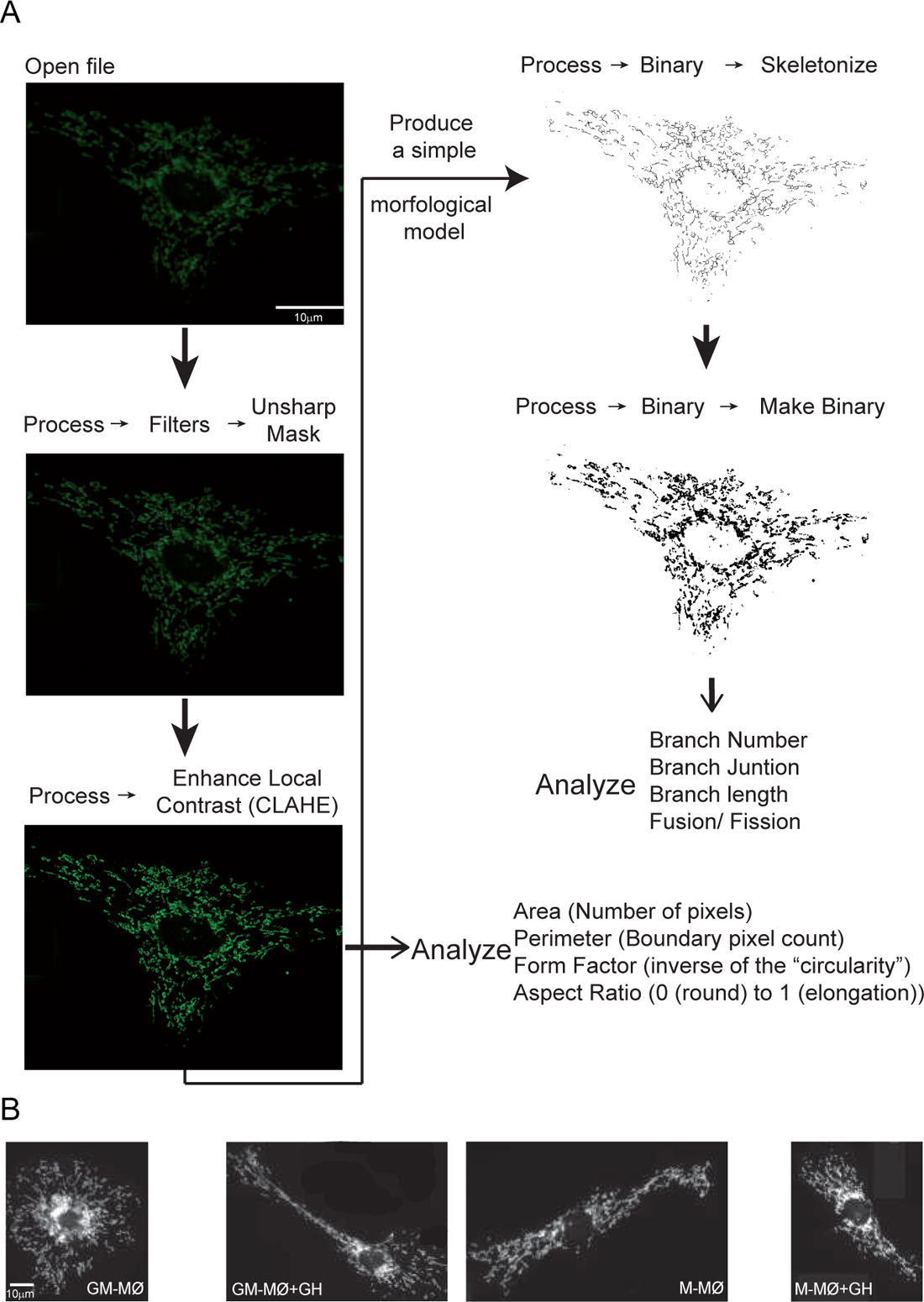

**Supplementary 3.**
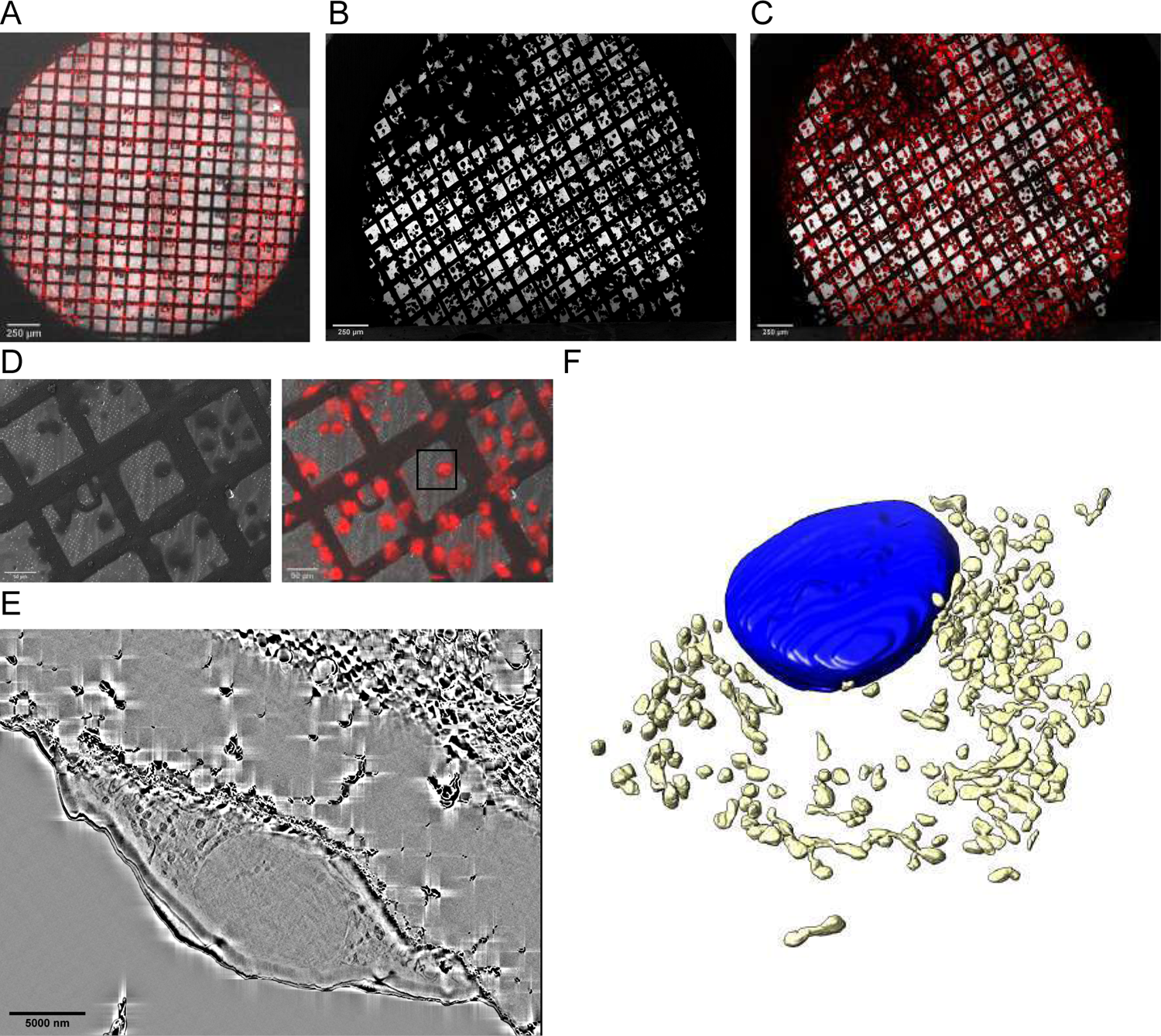

**Supplementary 4.**
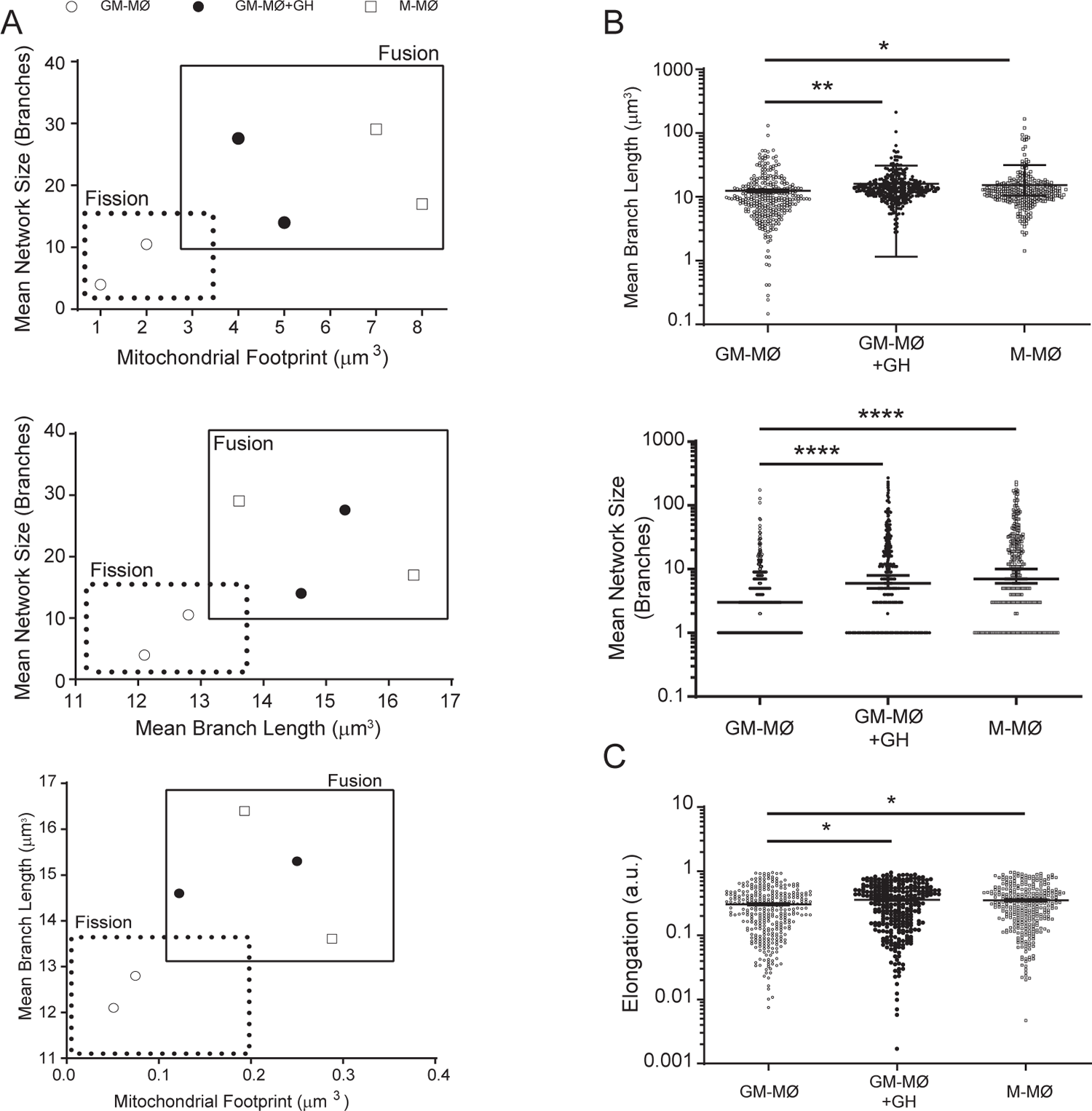

