## Supplementary Files for "Growth hormone remodels the 3D-structure of the mitochondria of inflammatory macrophages and promotes metabolic reprogramming"

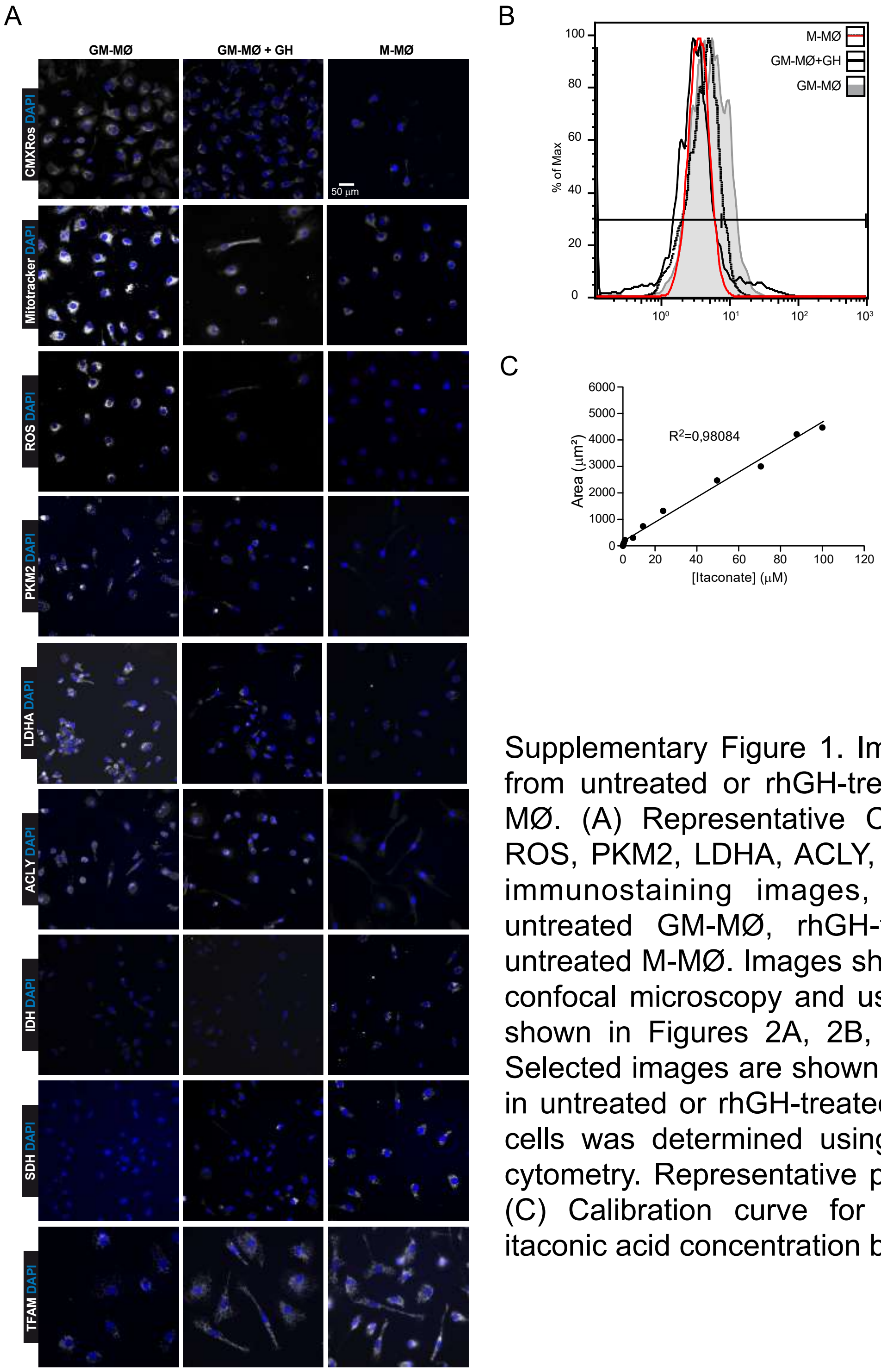


Supplementary Figure 1. Immunostaining images from untreated or rhGH-treated GM-MØ and M-MØ. (A) Representative CMXRos, MitoTracker, ROS, PKM2, LDHA, ACLY, IDH, SDH and TFAM immunostaining images, as indicated, from untreated GM-MØ, rhGH-treated GM-MØ and untreated M-MØ. Images shown were obtained by confocal microscopy and used to obtain the data shown in Figures 2A, 2B, 2E, 3A, 3D and 4C. Selected images are shown (n=8). (B) ROS levels in untreated or rhGH-treated GM-MØ, and M-MØ cells was determined using DCFH-DA and flow cytometry. Representative plots are shown (n=8). (C) Calibration curve for the determination of itaconic acid concentration by UV detection

Supplementary 2

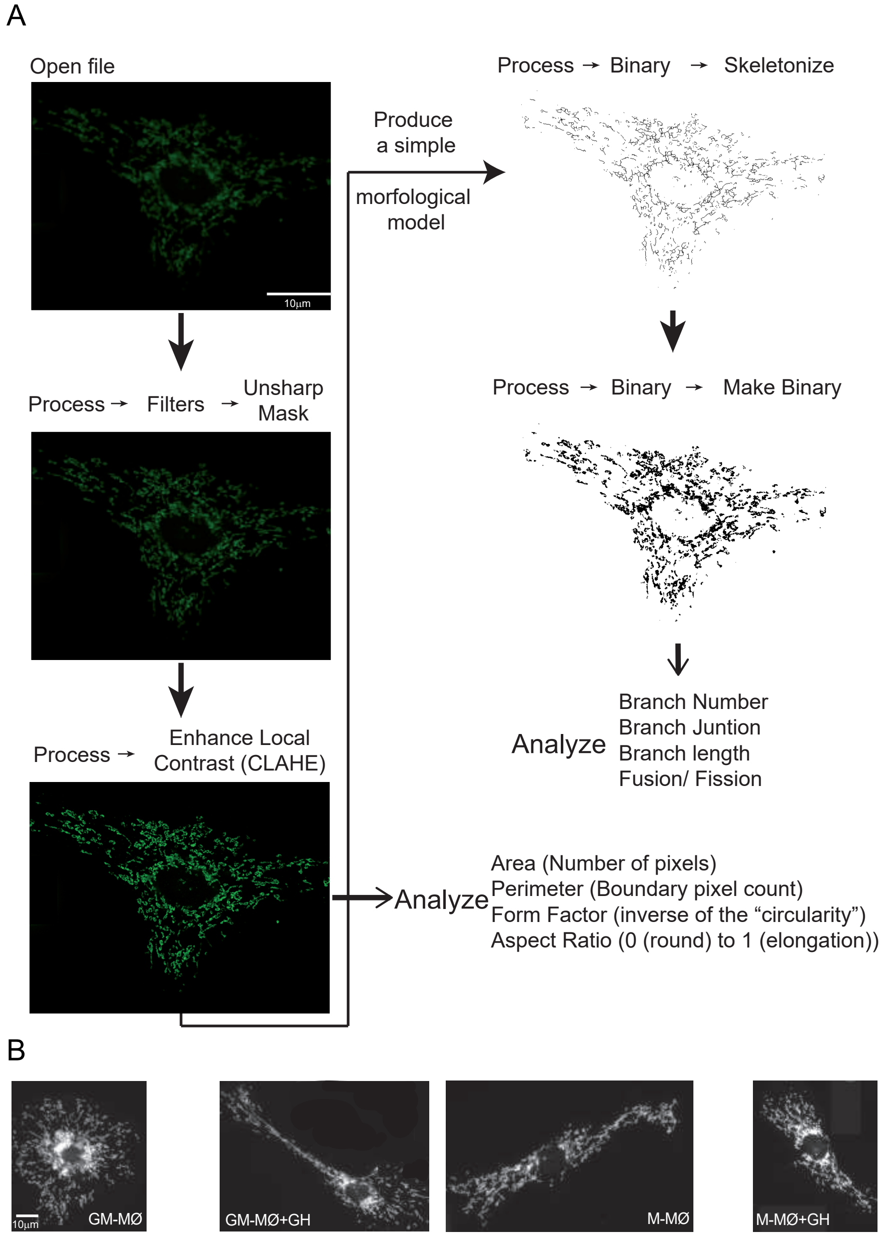


Supplementary Figure 2. Workflow of 2D images. (A). Workflow of 2D confocal microscopy images for the analysis of anti-TOM22-labeled samples. (B). Representative images of untreated or rhGH-treated GM-MØ and M-MØ stained with anti-TOM22 monoclonal antibody.

Supplementary 3

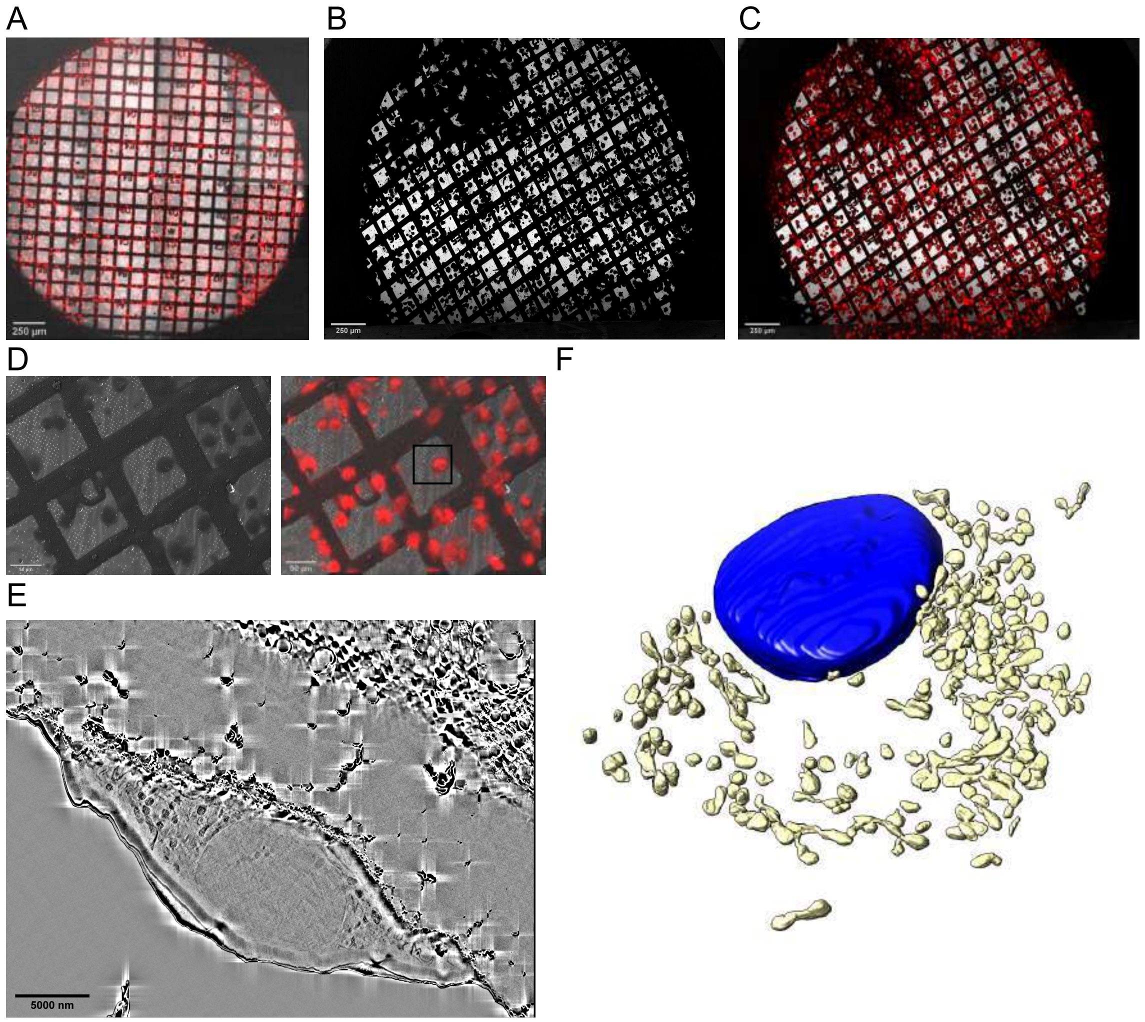


Supplementary Figure 3. Workflow of Cryo-FIB-SEM of a macrophages grown on a grid. (A). Cryo-confocal overlayed image of bright field and MitoTracker Red FM signal on a representative grid. Scale bar: 250 µm. (B). SEM image of the same sample shown in A. (C). Overlaying of cryo-fluorescence MitoTracker signal with SEM image shown in B. (D). On the left, SEM image of the macrophage chosen for analysis based on correlative information shown in C and on the right, same area highlighted (black square) in the correlated image. Scale bar: 50 µm. (E). A slice through a macrophage whole FIB-SEM volume after image processing. Scale bar: 5000 nm. (F). Three-dimensional rendering of the segmentation performed in the FIB-SEM volume shown in E (nucleus, blue; mitochondria, light yellow).

Supplementary 4

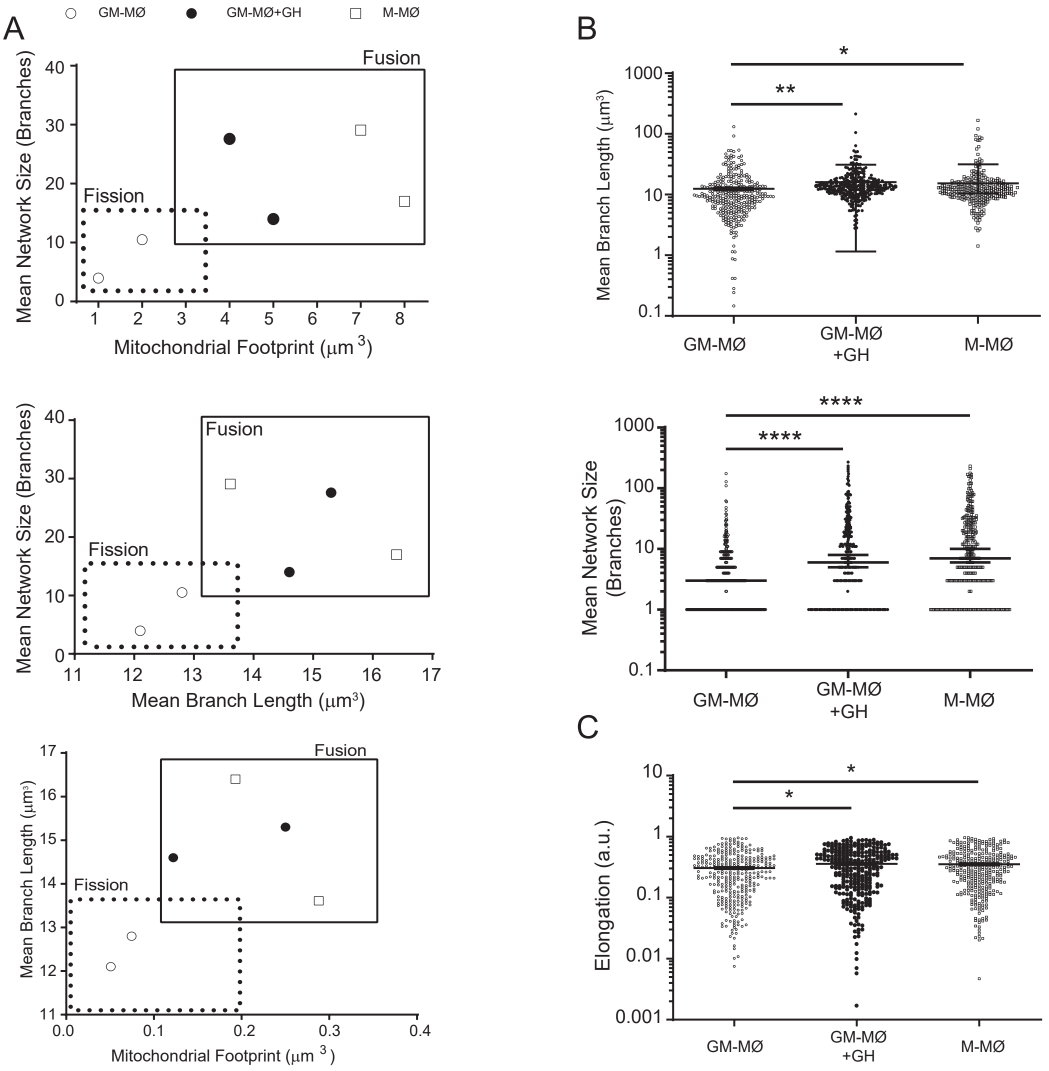


Supplementary Figure 4. Treatment with rhGH of GM-MØ modulates mitochondrial ultrastructure. (A). Two-dimensional scatter plots from mitochondrial network analysis (volume, network size, and branch length) comparing untreated or rhGH-treated GM-MØ, and M-MØ. The quadrants of fission/fusion were added to enclose clustered macrophages in feature space based on their mitochondrial organization. Dots are data of individual cells (n=3). All data shown are representative of three independent runs of experiments. (B). Quantitative analysis of network size and branch length of mitochondria from untreated or rhGH-treated GM-MØ, and M-MØ. Dots are data from individual cells (n=3) of three independent experiments. Results are expressed as mean ± SD. Paired t-test $*p<0.05$, $**p<0.01$, $***p<0.001$, $****p<0.0001$. (C). Quantification of the elongation of untreated or rhGH-treated GM-MØ and M-MØ. Statistical analysis as in (B).
